# Application of Whole Proteome Thermal Shift Assays to Define PERK-dependent Changes in Protein Homeostasis during the Unfolded Protein Response

**DOI:** 10.1101/2025.04.08.647882

**Authors:** Neil A. McCracken, Sarah A. Peck Justice, Whitney R. Smith-Kinnaman, HR Sagara Wijeratne, Avery M. Runnebohm, Stephane Pelletier, Emma H. Doud, Ronald C. Wek, Amber L. Mosley

## Abstract

The Unfolded Protein Response (UPR) is a cellular pathway activated by sensory proteins, including the protein kinase PERK (EIF2AK3), that monitors perturbations in the endoplasmic reticulum (ER). Using tunicamycin, which induces ER stress by thwarting N-glycosylation, we monitored system-wide changes in the proteome using PISA (Proteome Integral Solubility Alteration) and abundance analysis. Global proteomics revealed precise changes in membrane- and ER-associated proteins through widespread induction of ER-associated degradation (ERAD) while normalized PISA (nPISA) analyses selectively identified pathway changes associated with drug mechanism of action. nPISA analysis following tunicamycin treatment in cells, in combination with genetic disruption of PERK, facilitated identification of novel proteins involved in PERK-dependent and -independent processes and how those changes intersect with PERK function specifically during the ER stress response. Overall, protein-centered multiomics analyses defined the precise proteome alterations in tunicamycin-induced ER stress, highlighting the consequences of PERK disruption on ER-mitochondrial homeostasis.

## Introduction

Protein thermal shift analysis (TSA) has been used as a tool to monitor protein engagement with small molecules^1^. In recent years, these efforts have been expanded to facilitate small molecule target identification in system-wide analyses using quantitative proteomics methods including cellular thermal shift assays (CETSA), thermal proteome profiling (TPP), and Proteome Integral Solubility Alteration (PISA)^1–3^. Early system-wide CETSA studies suggested that both primary and secondary drug targets could be detected using these methods^2^. However, it has not been established if system-wide TSA methods can discriminate between small molecule induced changes in models with other perturbations such as genetic disruption. Addressing these key questions is important for understanding the utility of TSA-based methods for small molecule screening and in diverse cellular models with varied genetic backgrounds.

Maintenance of protein homeostasis (proteostasis) is vital for the function and survival of cells. In specialized secretory cells, levels of protein secretion can reach 800,000 proteins per minute^4^. To meet this demand, the Unfolded Protein Response (UPR) manages the critical balance between capacity and fidelity as proteins are synthesized, modified, and folded in the endoplasmic reticulum (ER) for subsequent passage through the secretory pathway. Induction of the UPR involves three ER transmembrane sensory proteins, PERK, IRE1, and ATF6 that function to sense accumulating unfolded proteins and trigger expression of genes that help expand the processing capacity of the ER and restore proteostasis^5,6^.

In the UPR, PERK is central for lowering global protein synthesis through induced phosphorylation of eIF2, which deters delivery of initiator tRNAs to the translation machinery. Consequently, there would be lowered influx of nascent proteins that would further challenge the protein processing apparatus in a stressed ER. Phosphorylation of eIF2 by PERK also leads to preferential translation of select gene transcripts that can serve critical adaptive functions during ER stress^7^. Among these adaptive genes is ATF4, whose enhanced expression leads to transcription of UPR-targeted genes that can serve to restore proteostasis^7,8^. In addition to PERK-directed translational control, other protein quality control pathways are induced by the UPR, including ER-associated degradation (ERAD) that targets problematic secretory proteins for decay. Chronic induction or dysregulation of these pathways is directly implicated in numerous human diseases including neurodegeneration^9^, cardiovascular disease^10^, cancer^11^, autoimmunity^12^, and diabetes^13^.

Proteomics approaches have been applied to cells undergoing ER stress to determine changes in nascent protein production to reveal the functional translatome and to probe alterations in protein structure through analysis of cysteine accessibility^14,15^. However, a comprehensive analysis of proteins with changes in biophysical state during ER stress has not been reported. In previous work, we showed that TPP can robustly identify the protein targets of small molecule inducers (SMI) of the UPR with short incubation times that allow for SMI-target interaction^16^. With ER stress induction, longer time points following SMI treatment are likely to be confounded by secondary signaling events such as alterations in gene expression and protein phosphorylation, along with changes in protein folding and turnover, which have not yet been resolved system wide.

This study pursues a comprehensive analysis of proteins with changes in biophysical state during early induction of ER stress. Here we apply the thermal profiling method PISA^3^ to characterize the dynamic changes of the whole proteome during ER Stress induction with a paired analysis of global protein abundance. We further extend these studies by combining ER stress induction with genetic perturbation of PERK signaling. The combination of drug treatment and knockout of this ER stress sensor facilitates the multiomics definition of UPR changes that are dependent on tunicamycin and/or PERK, providing high-quality datasets for the proteomics, drug discovery, and cellular stress response research communities. Overall, these data can be used as resources for identification of novel proteins involved in PERK-dependent and - independent processes that are involved in the ER stress response.

## Results and Discussion

### Tunicamycin treatment induces UPR and ERAD gene expression programs

Tunicamycin inhibits N-glycosylation^17^ in the initial steps of glycoform assembly in the ER^18^, and thapsigargin triggers release of calcium from the ER that is suggested to impede proper protein assembly. Illustrating the UPR induction, treatment of HEK293A cells with either Tm or Tg triggered activation of PERK as viewed by autophosphorylation that slowed migration of PERK by immunoblot analyses (Fig. 1B). Upon induction, PERK phosphorylation of the α subunit of eIF2 coincided with enhanced levels of ATF4 and XBP1s, which occurs via IRE1 signaling (Fig. 1B). We optimized the concentration and treatment regimen for Tm such that it was equivalent or exceeded the induction of hallmark UPR markers induced by Tg^16,19^ (Fig. S1A).

**Figure 1:**
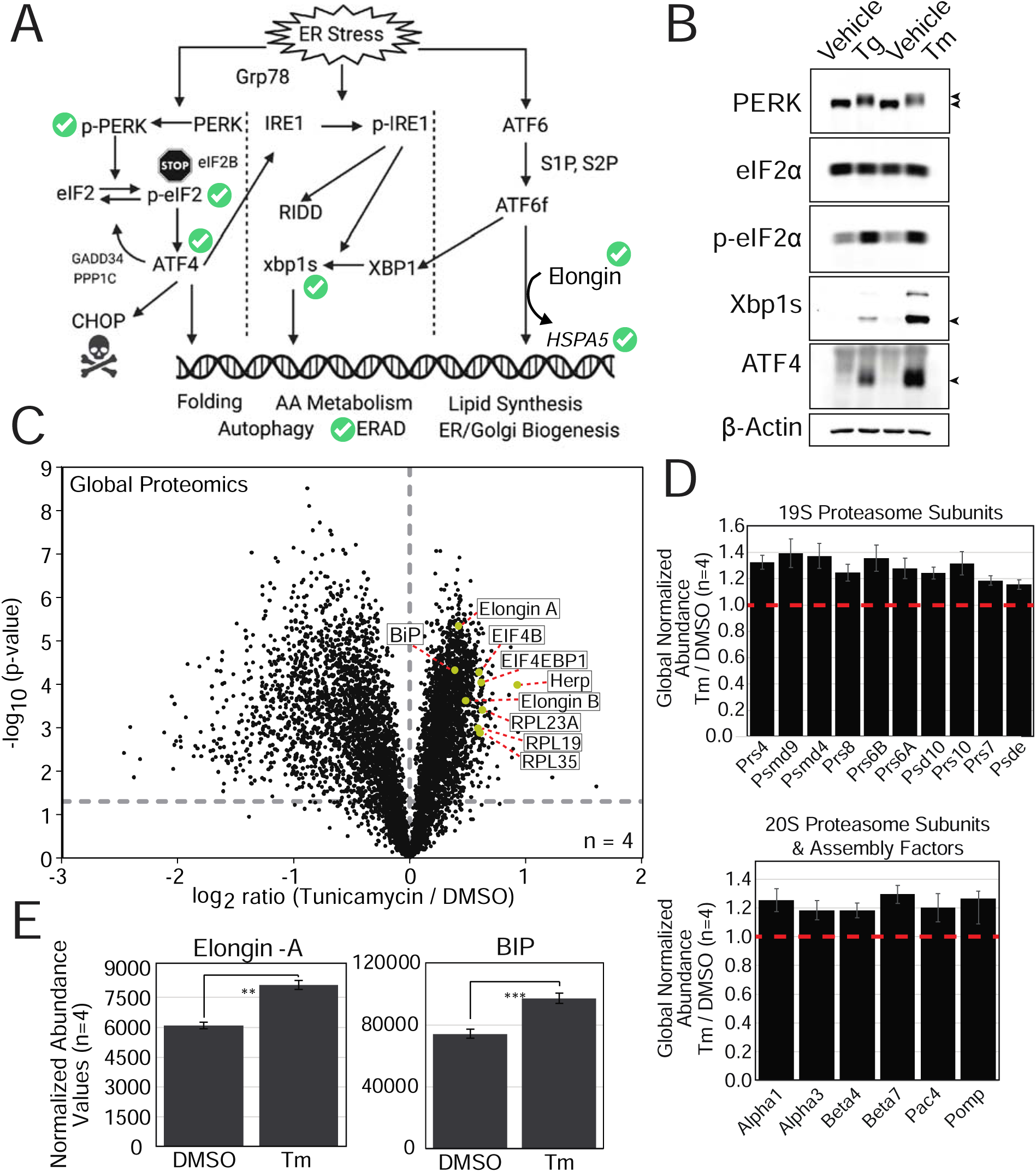
Tunicamycin treatment causes ER stress and wide-spread changes in the proteome. **A**. Schematic representation of ER stress signaling pathways induced by Tm treatment for 3hrs. Green checkmarks are shown for events that have been confirmed as active by global proteomics analysis. **B.** HEK293A cells were treated with 1 μM Thapsigargin (Tg) for 1 hour or 10 μM Tunicamycin (Tm) for 3hrs. Lysates were prepared and total and phosphorylated proteins were measured by immunoblot analyses. UPR markers in each blot are indicated next to each panel. β-actin was included as a loading control. **C**. HEK293A cells were treated with 10 μM Tm for 3hrs or 0.5% DMSO and global proteomics quantitation was performed (n=4). Volcano plot shows log_2_ ratio of abundance in Tm/DMSO for 6,181 proteins (black circles). Proteins of note are highlighted in yellow circles with corresponding labels. The gray dashed line designates a p-value cutoff of ≤0.05. **D.** Quantitative data showing increased abundance of subunits of the19S proteasome (top) and 20S proteasome (bottom). The red line indicates no change in abundance between Tm and DMSO. Data is shown as average ratios +/- standard deviation (n=4). **E**. Normalized abundance values for ATF6 transcription regulator Elongin-A and the protein product of the *HSPA5* gene BiP. Data is shown as average normalized abundance values +/- standard deviation with single replicate values shown as a dot (n=4).

We performed tandem mass tag (TMT) based global quantitative proteomics to profile the protein abundance changes following 10 μM Tm treatment (hence Tm treatment) or vehicle (hence DMSO) for 3 hours (Fig. S2, Fig. 1C, Table S1). Global proteomics analysis showed that this regimen of Tm treatment induced widespread changes in the abundance of steady-state proteins during ER stress (Fig. 1C). For example, Tm led to increased levels of HERP (Fig. 1C, in yellow), which is critical for ERAD and is reported to be induced by the UPR^20,21,22^. Further pathway analyses indicated enrichment in upregulated proteins involved in protein synthesis and SUMO-related signaling (Fig. S1B). Included among the upregulated proteins involved in protein synthesis were EIF4B and EIF4EBP1 (4EBP1) that regulate the initiation phase, along with large ribosomal subunits RPL19, RPL23A, and RPL35 (Fig. 1C, Fig. S1C). Expression of 4EBP1 has previously been shown to be induced through ATF4 signaling^23^. Numerous protein subunits of the 26S proteasome complex were also significantly upregulated when normalized to total protein abundance or β-Actin levels (Fig. 1D & E, Fig. S1D & E). These data agree with prior reports of proteasome transcript upregulation during UPR related processes including ERAD^24^. Proteasome assembly factors PAC4 and POMP were also significantly increased, further suggesting ERAD induction as previously described^24–26^. Global protein-level changes that reflect other UPR gene expression signatures include upregulation of the -A and -B subunits of Elongin, which aid in gene induction at ATF6 regulated genes including the *HSPA5* gene (encoding BiP)^27^. Protein levels of the key ER chaperone BiP were also significantly increased with Tm treatment (Fig. 1E, right).

Hierarchical clustering analysis of the top changes in global proteomics with UPR induction from Tm treatment as compared to DMSO treatment showed widespread protein abundance decreases across all biological replicates (n=4, Fig. 2A, adjusted p-value ≤0.05). The 530 decreased proteins were analyzed using ShinyGO for enrichment in Cellular Component terms, revealing enrichment in proteins resident to the ER, ER membrane, and the outer nuclear membrane (Fig. 2B). Proteins resident to the Golgi membrane were also statistically enriched (Fig. 2B). All together, these findings indicate that three major arms of ER stress signaling have been induced (Fig. 1A) through multiple gene expression and feedback loops upon 3hrs of treatment with tunicamycin with many protein decreases observed within the endomembrane system.

**Figure 2:**
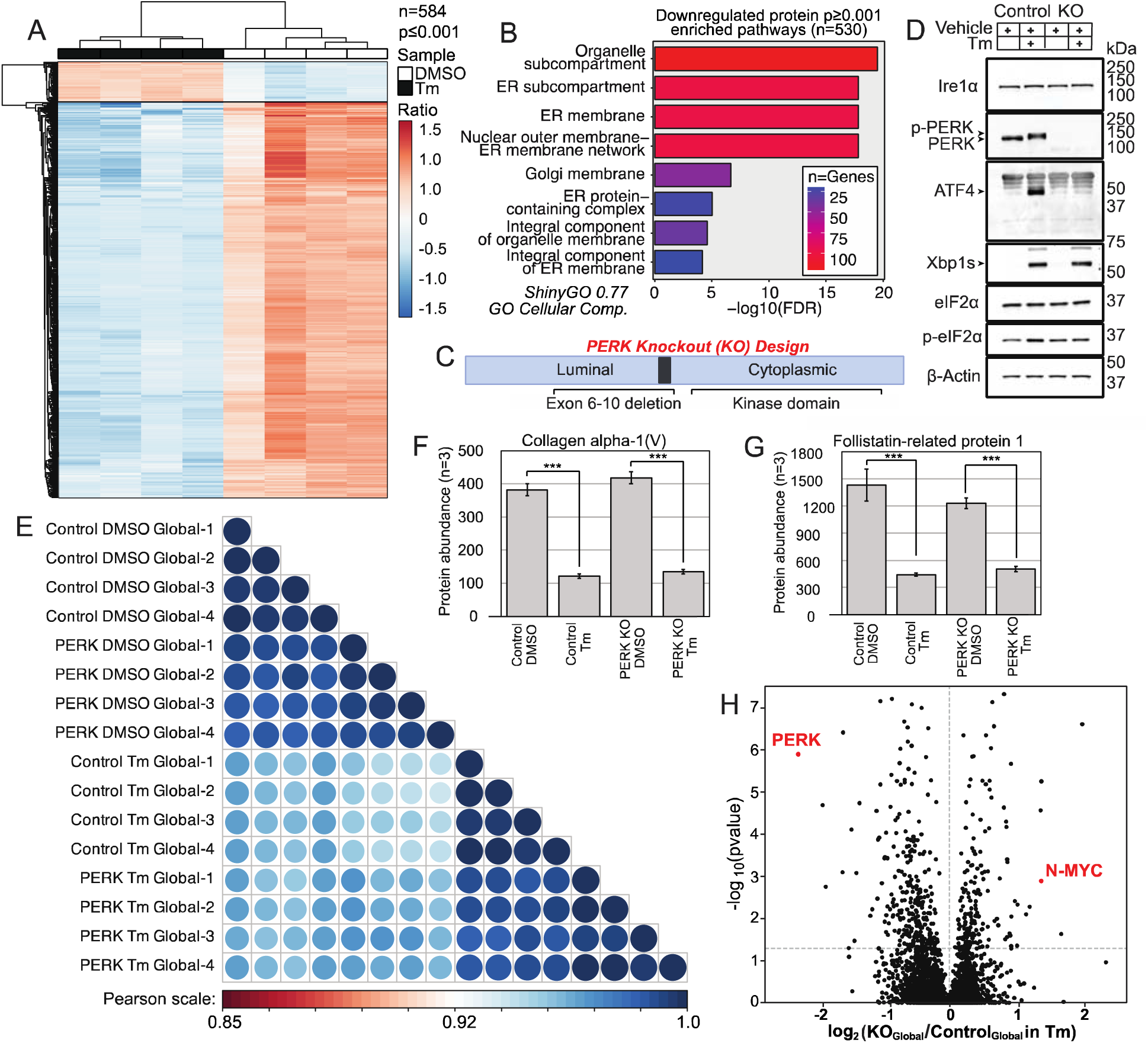
Global proteome changes observed with Tm treatment for 3hrs are not dependent on PERK. **A**. Hierarchical clustering analysis of global proteome changes with an adjusted p-value ≤ 0.001 (n=584 proteins). Biological replicates are shown as columns with each row representing values for a single protein. Tm replicates group to the left (black boxes) and DMSO replicates group to the right (white boxes). For analysis, pareto scaling was performed and the scale is shown to the right. **B**. ShinyGO analysis of Cellular Component with each ontology given to the right. Bars represent - log_10_ of FDR and bar color represents number of proteins. **C**. Schematic indicating location of exons 6-10 that were deleted by CRISPR/Cas9 targeting to generate a PERK KO cell line. **D**. Immunoblot analyses of markers of UPR induction; loading control was β-Actin. **E**. Correlation matrix comparing biological replicates (n=4) for global proteomics studies for PERK KO and WT clones treated with either DMSO or Tm. Pearson scale shows the color scale for the correlation R^2^ values. **F&G**. Normalized abundance values for (F) Collagen alpha-I (V) and (G) Follistatin-related protein I measured by global proteomics (n=3). Data is shown as average +/- standard deviation. **H**. Volcano plot showing changes in protein abundance in PERK KO cells / WT cells treated with Tm (n=4). The y-axis shows the -log_10_p-value for each protein measurement (represented by a dot). Red dots are shown for quantitation of PERK and N-myc. The dashed line on the figure represents a p-value cutoff of 0.05.

### Early UPR proteostasis changes alter ER-localized proteins and are largely independent of PERK

To delineate the contribution of PERK and translational control to the widespread changes observed in the global proteomics data, we produced PERK KO HEK293A cells by CRISPR-Cas9 deletion of exons 6-10 of the gene (Fig. 2C). Single clones were isolated and expanded for these studies and PERK KO was confirmed by DNA sequencing and immunoblot analysis of multiple clones (Fig. 2D, Figs. S3A & B). The control cells for these studies also underwent single cell clonal selection but retain wild-type (WT) PERK. Immunoblot analysis of Tm treatment samples confirmed that PERK protein was not detectable in PERK KO clones and that ATF4 protein induction did not occur following tunicamycin treatment in the absence of PERK (Fig. 2D). As expected, while levels of total eIF2α did not change with PERK KO, phosphorylation of eIF2α was not induced following tunicamycin treatment in KO cells compared to WT (p-eIF2α, Fig. 2D). PERK KO and WT cells contain a stably integrated P(AAREx6)-Luciferase reporter (Fig. S3C, described in ^28^) that was used to measure ATF4 transcriptional activity in a luciferase assay. PERK KO cells showed no significant induction of ATF4 activity following tunicamycin treatment, although WT cells showed significant induction (Fig. S3D). Phenotypically, PERK KO cells also did not show any major morphology differences or significant growth defects compared with WT in four days of growth as measured by an Incucyte cell density analysis (Fig. S3E, n=3). Together, these results support the utility of the PERK KO cells to study the contributions of this UPR branch on proteostasis.

To characterize proteostasis changes, we performed global proteomics analysis of PERK KO vs WT HEK293A cells in the presence or absence of Tm treatment for 3hrs through independent mass spectrometry analyses of Tm or DMSO treatment using TMTPro multiplexes (Table S2). This genotype-based analysis revealed fewer overall changes across the 7400 proteins quantified, suggesting a limited dependence of the UPR response on PERK at this early UPR time point (Fig. 2D). Pearson correlation-based analysis was performed on the Tm and DMSO datasets comparing PERK KO to WT clones with a focus on proteins quantified in both conditions. Correlation analysis confirmed that Tm or DMSO treatment was the major driver of differences between the four datasets, clearly showing that the widespread decreases caused by Tm treatment are not dependent on PERK translational control, but rather likely depend on a protein degradation process such as ERAD (Fig. 2D). For example, significant abundance changes were observed for two secretory proteins which travel through the endomembrane system for eventual secretion: COL5A1 and FSTL1 (Fig. 2F & G). Both COL5A1 and FSTL1 show rapid and reproducible decreases in abundance following Tm treatment in both genotypes. Together, these findings clearly illustrate that decreases in protein abundance associated with Tm treatment at 3hrs are mostly PERK-independent.

Although there were few protein abundance differences with >2-fold change identified between PERK KO vs WT HEK293A cells following 10 μM Tm treatment for 3hrs, there are targets that display clear PERK dependence (Fig. 2H, Fig. S3F, and Table S2). Major changes of note in the Tm treatment dataset include PERK itself, with an expected abundance decrease observed in PERK KO cells, and N-Myc which is significantly upregulated in PERK KO cells from Tm treatment (Fig. 2H, red) but not in DMSO (Fig. S3F, red). N-Myc has a half-life that ranges from 30-50 minutes in previously published studies^29,30^. In the absence of PERK, Tm does not cause decreased protein synthesis and there is continued production of proteins with rapid turnover such as N-Myc resulting in their accumulation relative to WT cells (Fig. 2H). In PERK KO vs. WT with DMSO, N-Myc levels did not significantly change (Fig. S3E). Together, these findings suggest that 3hrs of Tm SMI treatment is sufficient for induction of UPR- and ERAD-related proteins and widespread decreases in ER and endomembrane system proteins. Although proteome changes associated with PERK signaling, including induction of ATF4 and its downstream target 4EBP1, were observed with 3hrs of Tm treatment, the majority of proteome changes were PERK-independent, suggesting that protein reduction at this time point is largely achieved through clearance mechanisms such as proteasome-dependent ERAD.

### Analysis of the UPR by thermal profiling analysis using PISA

Although global proteomics analysis gave multiple insights into proteostatic changes that occur following 3hrs of Tm treatment, the method did not provide specific insights into the MOA and downstream consequences of small molecule treatment. While it has been established that TPP-based approaches can identify small molecule target proteins^31^, few of these studies have sought to identify MOA changes in signaling conditions with broad protein biophysical changes such as those in UPR. We applied a TMTpro-based LC-MS/MS PISA assay as initially reported by Gaetani et al^3^ to measure changes in protein biophysical status. In prior work, we determined that the average melt temperature for proteins in HEK293A cells was close to 50°C^32^, in agreement with prior meltome studies^33^. The heat gradient applied for PISA experiments included 8 distinct temperatures from 40-60°C to capture the melting temperature range for most human proteins^33^. A major adaptation that was made for this study was to normalize PISA values by the global proteome abundance measurements to calculate normalized PISA abundance changes (nPISA)^3^. This objective was facilitated by splitting the same biological samples for both global and heat treatment for PISA analysis in a multiplex design where global and PISA biological replicate samples (n=4) were analyzed as a single 16-plex (Fig. S2). These steps minimized biological and technical variability, in addition to eliminating missing quantitative measurements needed for normalization. In traditional PISA analysis, an observed increase in protein abundance with the same magnitude of the global change could lead to the false conclusion that protein stability is affected by treatment as we observed for the UPR-induced ERAD protein HERP (Fig. S4A) and for BiP. The nPISA values account for the influence of UPR-induced gene / protein expression or protein degradation changes to limit misinterpretation of biophysical state changes (Fig. S4B).

Since Tm disrupts N-glycan synthesis through inhibition of UDP-GlcNAc production, an early step in protein N-glycosylation, we highlight the enzyme groups involved in UDP-GlcNAc production and downstream processes in N-linked glycosylation in the ER without our nPISA dataset (Fig. 3A & B, enzyme group designated with numbers indicated in Fig. 3B). Group 1 proteins include the dolichyl-phosphate Glc-NAc-1-phosphate transferase (DPAGT1) which is the known target of tunicamycin. While the global abundance of DPAGT1 did not change significantly between DMSO and drug (Table S1), there was a statistically significant increase in DPAGT1 nPISA levels with Tm treatment indicating that drug binding stabilizes DPAGT1 (Fig. 3C). The stabilization of many drug targets from drug engagement has been observed previously in other CETSA, TPP, and PISA studies, but has not previously been shown in a complex proteome background with large protein abundance changes such as in early UPR signaling^32,34,35^.

**Figure 3:**
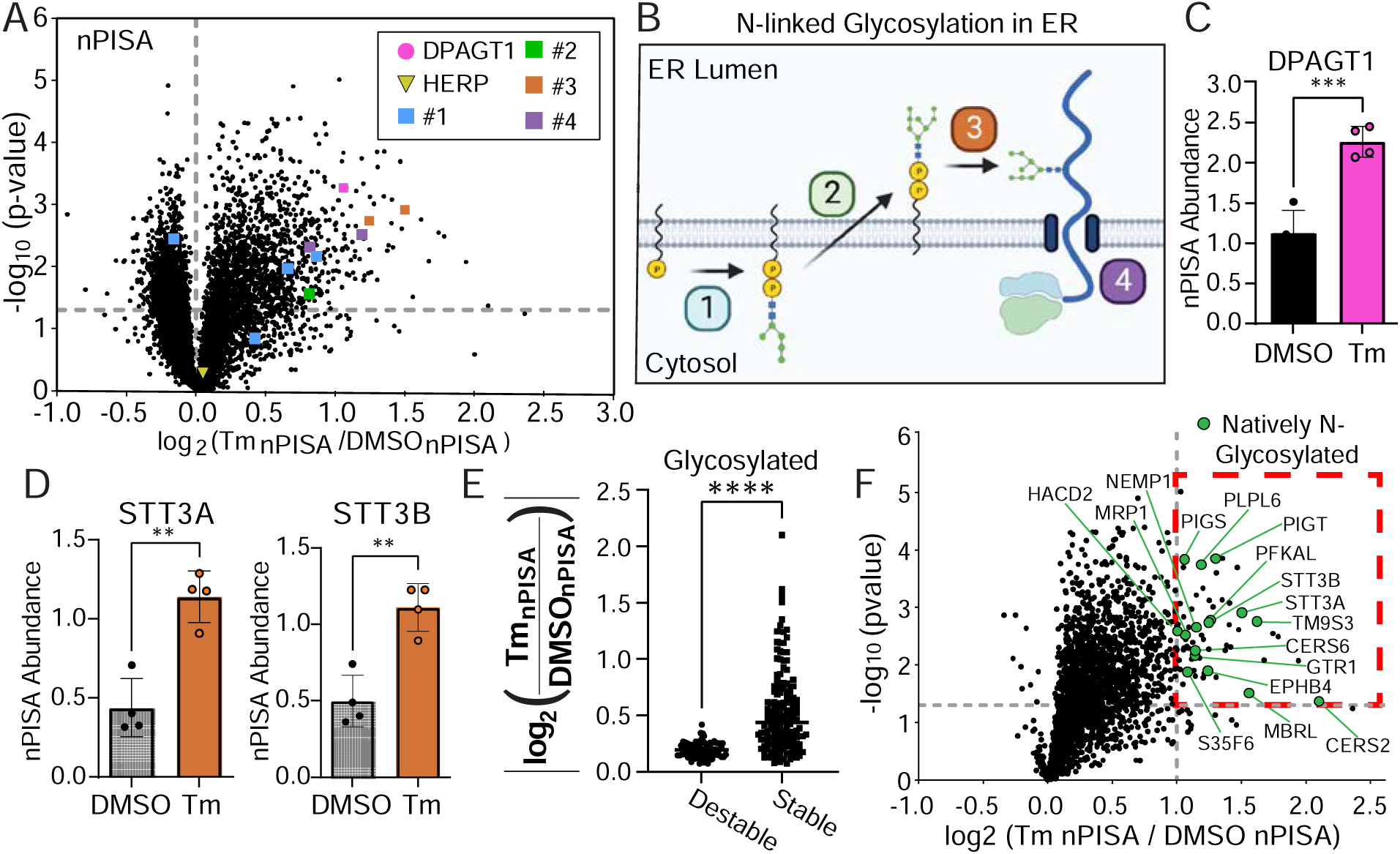
Normalized PISA (nPISA) analyses identify changes following Tm treatment. **A**. Volcano plot of nPISA data from HEK293A parental cells treated with Tm or DMSO (n=4). Pink circle is DPAGT1; the yellow circle is HERP. Protein groups 1-4 are highlighted based on N-glycosylation functional pathways outlined in B. **B**. Model representing key enzyme groups associated with N-glycosylation: 1. UDP-Glycan production. 2. Glycan flipping into the ER Lumen. 3. Glycan transfer onto the nascent polypeptide. 4. Polypeptide translocation into the ER Lumen. **C&D**. Average nPISA abundance values for (C) Tm target protein DPAGT1 and (D) oligosaccharyl transferase (OST) complex subunits with values for replicates indicated with dots (n=4). **E.** Absolute value plot showing the data distribution for natively glycosylated proteins (Uniprot) nPISA measurement. Each dot represents an average nPISA value for one protein. **F.** Volcano plot of proteins that were significantly decreased in abundance but stabilized according to nPISA values at least 2-fold (log2 nPISA ≥ 1; dashed red box).

Considering the biochemical and biophysical importance of N-linked protein glycosylation (hence N-glycosylation) we anticipated that inhibition of DPAGT1 would have extensive consequences on downstream membrane protein processing. UDP-Glycans produced by group 1 enzymes (designated in Fig. 3A in blue) are flipped from the cytosolic face of the ER to the ER lumen for further maturation by group 2 (green, Fig. 3B) enzyme ALG9, which was also found to be thermally stabilized following tunicamycin treatment (Fig. 3A & B, Fig. S4C, Table S1). Group 3 enzymes (orange, Fig. 3B) STT3A and STT3B are subunits of the oligosaccharyl transferase (OST) complex and show a similar magnitude of response to tunicamycin treatment, consistent with their protein-protein interactions and similar functional roles in glycosyltransfer (Fig. 3D, Table S1). Finally, proteins that are subjected to N-glycosylation are translocated into the ER lumen by the group 4 enzyme SEC61 complex, which is also thermally stabilized in our nPISA analysis (Fig. 3A, S4D in purple). These nPISA studies provide evidence that both the target of tunicamycin DPAGT1 and downstream enzymes in the N-glycosylation pathway including transferases and translocons are significantly stabilized by Tm treatment. Gene Ontology (GO) analysis confirms a significant pathway enrichment in N-glycan biosynthesis specifically in the nPISA dataset when considering all proteins with adjusted p-values ≤ 0.05 (Fig. S4E). Additional enrichment was found for ABC transporters and steroid biosynthesis, which are both processes that involve multiple membrane proteins and in many cases N-glycosylation^36–39^. These findings show that nPISA analysis can effectively identify the target of Tm, DPAGT1, while selectively identifying additional protein thermal stability changes associated with Tm MOA.

It is expected that tunicamycin treatment leads to wide-spread defects in N-glycosylation leading to UPR induction. We found that a larger number of natively glycosylated proteins are thermally stabilized rather than destabilized based on nPISA, and in multiple cases there was a larger magnitude of biophysical state change (Fig. 3E). In addition, we found an enrichment in nPISA stabilization for proteins that have been identified as HSP targets in a prior study (Fig. S4F, ^40^) and proteins that contain coiled-coil domains (Fig. S4G). Prior work has established that the induction of the UPR allows for BiP-dependent clearance of both unfolded proteins and some protein aggregates, although proteins that cannot be correctly processed / folded are often subjected to ERAD^41–44^, as may be directly indicated by our global proteomics findings. To further analyze the population of proteins that were significantly stabilized by nPISA, we focused on proteins that were significantly decreased in global abundance but had ≥ 2-fold increases in thermal stability by PISA (Fig. 3F). In this strong responder group, we identified many proteins that are known to be natively modified by N-glycosylation consistent with cellular treatment with tunicamycin (Fig. 3F, natively glycosylated proteins indicated in green per Uniprot annotations^45^). Furthermore, most strong responders were membrane proteins with the most significantly enriched fraction of stabilized proteins being ER membrane proteins (Fig. S4H). Overall, proteins that participate in N-glycan biosynthesis, target proteins that are post-translationally N-glycosylated, membrane proteins, and proteins that required HSPs for folding were thermally altered following Tm treatment. *In-silico* thermodynamic prediction of protein free energy state is anticipated to be lower for glycosylated proteins^46^. Yet it has also been shown that proteins lacking proper glycosylation in the ER can associate with chaperones^47^ or aggregate^48^; scenarios that may increase the stability of these proteins, consistent with our results.

### PERK contributes to protein thermal stability changes during the UPR

PISA analysis was performed in PERK KO compared to WT cells to identify PERK-dependent changes in the response following treatment with Tm for 3hrs. In the case of the previous PISA experiment, the change in stability was measured between Tm treatment and DMSO (Fig. S2). For these nPISA experiments, the PISA and global samples were multiplexed so that the impact across the two cell lines (PERK KO / WT) could be directly measured in separate LC-MS/MS experiments (Fig. S5A). To allow for these comparisons, the first multiplex included the Global and PISA lysates from the Tm experiment while the second multiplex included lysates from DMSO treatment (Fig. S5A). At the 3hr time point, we observed a total of 202 proteins that were significantly stabilized in PERK KO cells and the vast majority of these (192/202) were only found to be stabilized in tunicamycin treatment (Fig. 4A). These findings strongly suggest that PERK is required even in early UPR to maintain homeostasis of specific protein targets.

**Figure 4:**
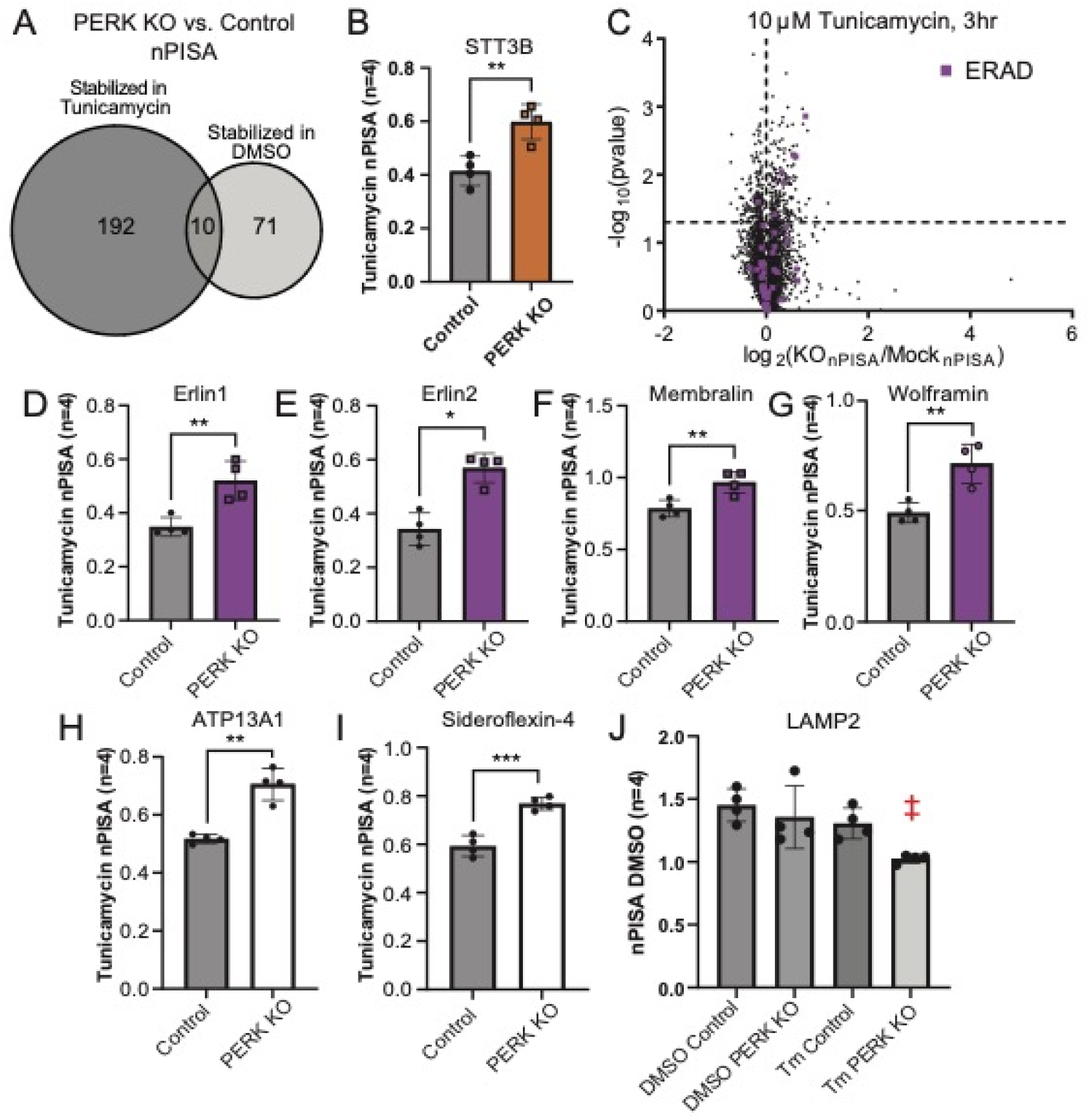
Loss of PERK leads to increased thermal stabilization proteins involved in ERAD and mitochondrial homeostasis. **A.** Venn diagram showing overlap between proteins that are thermally stabilized by PERK KO with Tm (left) or DMSO (right). **B**. STT3B nPISA values in Tm. nPISA values represent the average values plus or minus standard deviation (n=4). Individual biological replicate values are provided as dots in each bar. **C.** Volcano plot showing the data distribution for nPISA abundance values in PERK KO vs. WT cells. Each dot represents the average value for a single protein. Proteins that have been annotated to be involved in ERAD are designated by purple dots. Dashed lines indicate a p-value ≤ 0.05 on the y-axis and the 0 value on the x-axis. **D-I**. nPISA values for (D) Erlin1, (E) Erlin2, (F) Membralin, (G) Wolframin, (H) ATP13A, and (I) Sideroflexin-4. **J**. LAMP2 nPISA values provided as in panel B, with both DMSO and Tunicamycin treatment datasets are shown. The double cross indicates a two-way ANOVA analysis of genotype x Tm treatment (p-value ≤ 0.01).

STT3A and B were both significantly stabilized in parental HEK293A cells from tunicamycin treatment relative to the DMSO (Fig. 3D). We observed further exacerbation of OST subunits STT3A, STT3B, RPN1, and OST48 stabilization when PERK is lost (Fig. 4B and Table S2). Comparison of these findings to STT3A & STT3B stabilization in Fig. 3D, suggest that stabilization of the OST complex caused by Tm treatment is partially mitigated by PERK signaling. In both PISA datasets, we find that OST complex subunits are a highly responsive marker of Tm-induced UPR stability changes that are PERK dependent. It is possible that OST enzymes are being sequestered for degradation because of their proximity to ER translocases and hence ERAD machinery. The reproducible and significant stabilization of STT3A, STT3B, Sec61A and Sec61B are supported by engagement of these components with nascent proteins as they are co-translationally translocated into the ER lumen from the ribosome. It is also plausible that OST and SEC61 stabilization occurs related to protein synthesis “stalling”; a phenomenon that has been reported to occur during UPR to aid in degradation processes of nascent polypeptides^49^. Intriguingly, STT3B is a well-described major target of the immunoproteasome presented by MHC complexes and found in multiple immunopeptidomics studies. Our findings that OST subunits are highly sensitive markers of ER dysregulation by Tm in both WT and PERK KO studies suggests that OST protein stability could serve as a sensitive biomarker for ER stress, UPR, and/or ERAD.

Multiple other ER membrane proteins involved in protein processing showed increased thermal stability in PERK KO cells relative to WT, such as the signal peptidase complex (SPC) subunit SPCS3 that catalyzes signal sequence cleavage from nascent proteins as they are translocated into the ER (Table S2)^50^. DAVID pathway analysis revealed that the stabilized proteins in the Tm dataset that compared PERK KO to WT are highly enriched in transmembrane domain-containing proteins (109 total proteins, Benjamini FDR = 5.7 E-23) and ER membrane proteins (57 proteins, Benjamini FDR = 8.1 E-15). These findings reinforce the idea that PERK is required to reduce the impact of ER stress on membrane protein homeostasis (Fig. S5B & C). In the DMSO dataset, proteins found to be thermally stabilized in the absence of PERK also showed enrichment in transmembrane domain-containing proteins (31 total proteins, Benjamini FDR = 3.1 E-04) and ER membrane proteins (17 proteins, Benjamini FDR = 2.0 E-02) although for less targets than observed in Tm treatment (Fig. S6). The baseline stabilization of membrane proteins in the absence of PERK in DMSO treatment cells illustrates the importance of PERK for basal ER protein processing regulation (Fig. S5B&C compared to S6). These data can serve as a valuable resource for identification of membrane proteins with a strong dependence on PERK-related signaling for ER processing in both basal and UPR signaling states.

Upon loss of PERK, several targets were selectively stabilized relative to WT cells treated with Tm for 3hrs related to ERAD pathways (Fig. 4C). PERK-dependent thermal stabilization of ERAD-related proteins Erlin1 and Erlin2 was observed (Fig. 4D). The Erlins are ER-specific membrane proteins that are implicated in protein retrotranslocation from the ER lumen to the cytosol (Fig. 4D&E)^51^. The Erlins have also been shown to interact with ERAD machinery including the ERAD E3-ligases AMFR and RNF185^52–57^. Membralin, a previously described interaction partner of the Erlins^58^, also showed increased thermal stability by nPISA in PERK KO cells relative to WT in Tm treatment (Fig. 4F). The ER-mitochondrial membrane microdomains (MAMs) enriched protein Wolframin is also significantly thermally stabilized in PERK KO following Tm treatment (Fig. 4G)^59^. Wolframin and PERK have been shown to interact, to be highly enriched at MAMs, and to play a role in ER calcium homeostasis^59–61^. Wolframin has also been implicated in ERAD^62^.

In PERK KO cells, Tm treatment also led to thermal stabilization of several other proteins implicated in mitochondrial membrane protein homeostasis including ER-resident protein ATP13A1 (Fig. 4H) and multiple mitochondrial membrane proteins including sideroflexin-4 (Fig. 4I). Finally, we observed thermal destabilization of chaperone LAMP2A – a key protein involved in chaperone-mediated autophagy in PERK KO. LAMP2A levels have been reported to increase following Tm treatment^63^; however, our nPISA showed that thermal destabilization of LAMP2A only occurred in a PERK-dependent fashion (Fig. 4J, two-way ANOVA p-value = 0.0123). Together, these data show that PERK mitigates the deficiencies caused by the UPR such that PERK KO changes the thermal stability of numerous proteins involved in ERAD and ER-mitochondrial contact homeostasis.

### UPR signature analysis by quantitative proteomics-based MultiOMICs

A quantitative description of the UPR at the protein level remains elusive due to large scale protein changes including increased or decreased protein synthesis for specific targets. While some models do exist for describing the overall cellular changes during UPR, the transition from adaptive to terminal UPR is still not fully understood^64^. We reasoned that comparison of global proteomics and nPISA protein-based changes during Tm-induced UPR would allow for discovery of novel proteins that contribute to various UPR phenotypes without *a priori* knowledge. An overview of the significant changes in our datasets revealed that proteins segment into multiple functional groups with low interdependence (Fig. 5A). For example, Clusterin decreased in protein abundance and has decreased thermal stability by nPISA following Tm treatment. Clusterin is a known protein chaperone^65^ but most of its functions have been characterized as extracellular. Clusterin is known to also have cellular retained forms and has been shown to associate with BiP during ER Stress^66^; but it has not been functionally annotated in GO with ER Stress or related pathways.

**Figure 5:**
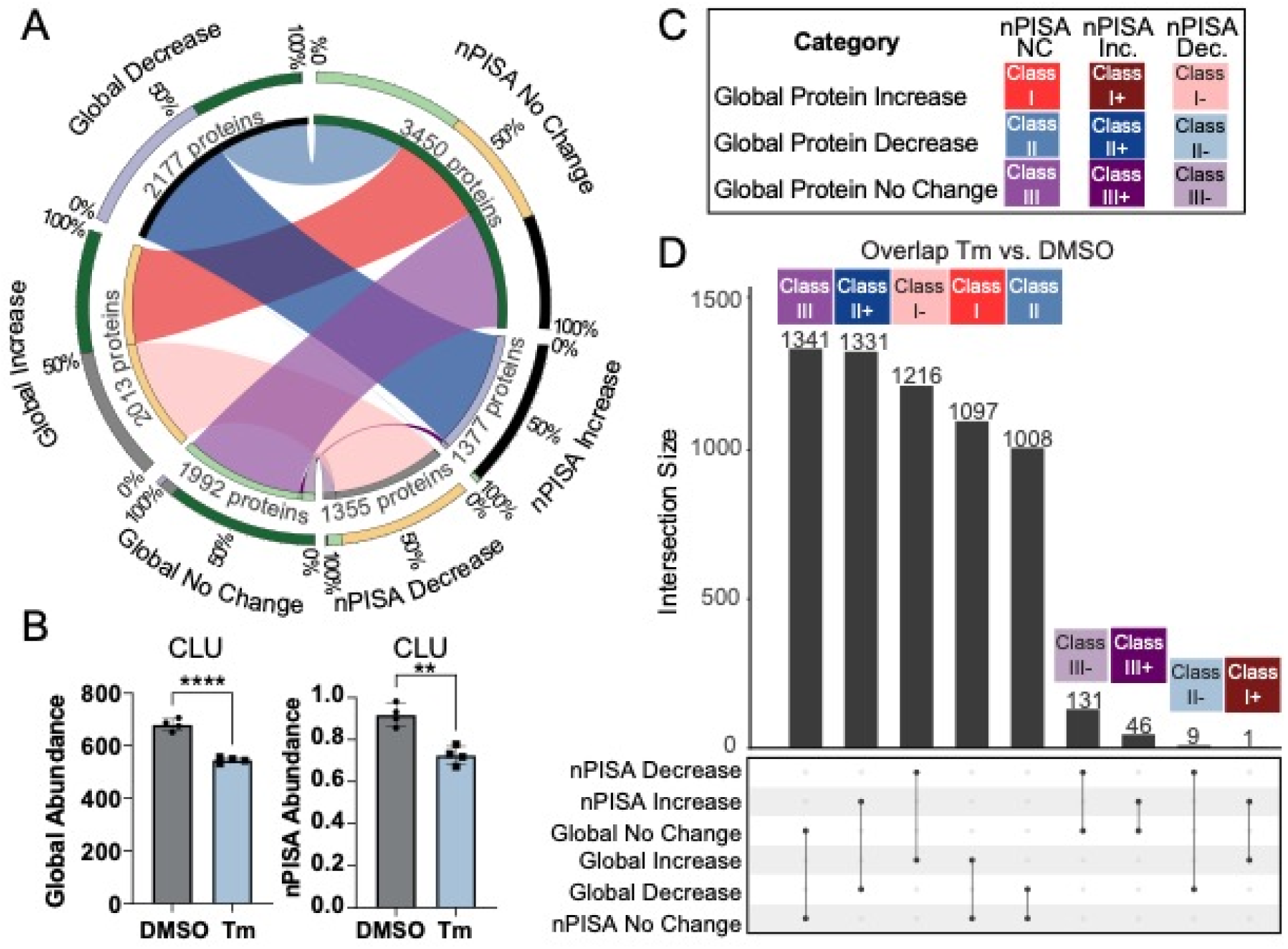
Discovery-based analysis of UPR classes through protein-based multiomics. **A.** Circos plot illustrating the quantitative proteomics subgroups for Global Proteomics or nPISA with changes defined as p-value ≤ 0.05. Ribbons within the plot indicate number of proteins that intersect between separate quantitative groups, color coded according to UPR Signature Class. The first tier defines the total number of proteins within each category. The outermost tier shows the percentage of each category that intersects with others. **B**. Clusterin (CLU) global abundance and nPISA values. Values represent average values +/-SD from Tm (blue bars) or DMSO (gray bars, n=4). Biological replicate values are provided as dots. **C**. Description of class definitions based on their intersections between defined quantitative groups. Class names are defined based on protein quantitation in global proteomics with a plus (+) or a minus (-) designating nPISA-based stabilization or destabilization, respectively. **D**. Upset plot illustrating the distribution of the different intersecting groups. Bar height indicates the total number of proteins within a specific class intersection (y-axis); the graph under the bar plot shows the intersecting groups (x-axis). Color-coded class categories are indicated in boxes above each bar on the plot. Table S3 provides a list of protein identities for each class.

To systematically provide intersection analysis of the global proteomics and nPISA datasets, we provide annotation independent functional subsetting of both characterized and uncharacterized proteins through multiOMICs classification (Fig. 5C & D). We chose to assign proteins to classes based on their global abundance status with Class I being all proteins that increase, Class II being all proteins that decrease, and Class III being proteins that did not have significant changes (Fig. 5C). Next, the thermal stability data was added to the classifications with significant increases in thermal stability indicated with a “+” and decreased thermal stability indicated with a “-”. This analysis results in 9 functional categories to define protein functional categories during UPR (Table S3).

The largest functional category identified was Class III with proteins that were not significantly changed in this study (Fig. 5D); however, many of our novel functional classes with significant changes included over 100 distinct proteins per class in 6 out of 9 categories. To illustrate the utility of these UPR functional classes, DAVID analysis was performed for each of the classes with results included to describe various functional insights provided in Fig. S7. HERP and various subunits of the 26S proteasome are defined as Class I proteins in the tunicamycin treatment dataset (Table S3). HERP has been reported to facilitate ERAD through ubiquitin-dependent proteasome degradation^67,68^ and is anticipated to increase in abundance with cellular stress signaling. Overall, protein chaperones are statistically enriched in Class I-by DAVID analysis with 66 total chaperones enriched associated with the Gene Ontology Molecular Function annotation = chaperone (Fig. S7 and Table S3). Multiple J-domain containing Hsp40 proteins, which are co-chaperones of BiP, were upregulated at the protein level and showed significant destabilization by nPISA^69^. Class I-protein DNAJB11 (also known as ERDJ3) is a well characterized ER lumen localized co-chaperone for BiP that facilitates ERAD of misfolded proteins^70–72^. DNAJB11 showed clear nPISA-based destabilization following tunicamycin treatment which could occur through engagement of unfolded clients, consistent with its roles defined in previous studies^73,74^. Small glutamine-rich tetratricopeptide repeat-containing protein alpha (SGTA) is a Class I-protein and co-chaperone that is not a member of the HSP40 co-chaperone family.^75^

The DAVID analysis for Class II proteins showed a relative decrease in abundance of proteins containing transmembrane domains, disulfide bonds, and/or N-glycosylation PTMs (Fig. S7). Class II is consistent with proteins that would be processed in the ER and may be degraded by ERAD during UPR signaling to prevent trafficking of improperly folded proteins. Although there are many Class II proteins that did not show changes in thermal stability (n=1008, Fig. 5D), there are many proteins that group into Class II+ that are decreased in global abundance but increased in thermal stability (n=1331, Fig. 5D). Although Class II+ is functionally enriched in transmembrane protein categories, Class II+ proteins additionally show functional enrichment as ER-specific membrane proteins with some related to protein processing and modification. This includes subunits of the SEC61 translocase and many other proteins that play a role in protein processing and folding in the ER. The chaperone DNAJC10, for instance, is one of a few BiP (GRP78) co-chaperones that is a member of Class II+^76^. DNAJC10 (also known as ERDJ5) contains both a J-domain and thioredoxin-like domains that have disulfide bond reductase activity. This additional functionality distinguishes DNAJC10 from many of the J-domain proteins enriched in Class I-. ERO1-like protein beta (ERO1B) is an additional protein involved in disulfide bond formation in the ER and a member of Class II+^77,78^. The stabilization of numerous ER functional classes suggests that increased protein interactions within the ER, possibly with the ERAD machinery as unfolded or improperly processed proteins, are transferred from the ER to the cytosol for degradation during UPR.

### UPR signature analysis of PERK KO cells identify changes in mitochondrial function and reduction-oxidation

Many of the protein abundance changes that were observed in Tm treatment compared to DMSO were found to be PERK-independent (Fig. 2). In class comparisons, we found that most proteins in the PERK KO datasets did not change in global abundance or thermal stability by nPISA (Fig. 6A and Fig. S8A). However, specific changes did occur that provide molecular insights into PERK contributions to adaptive UPR signaling (Fig. 1). Both PERK KO vs. WT datasets revealed significant changes in protein abundance and unlike changes by nPISA (Fig. 4A), there was substantial overlap between Tm and DMSO global abundance findings with 52.7% or 43.6% overlapping targets increasing (n = 314) or decreasing (n = 356), respectively, in both datasets (Table S4). These MultiOMICs data suggest that the abundance of many protein targets are regulated by PERK in basal conditions independent of ER Stress since changes were identified in both treatments. Analysis of the PERK-dependent proteins revealed that TIMMDC1, an assembly chaperone for the mitochondrial NADH: ubiquinone oxidoreductase complex (complex I) was significantly decreased by 40% in both datasets (p-values ≤ 0.05, Tables S2 & S4, Fig. 6B). There are also significant decreases in multiple subunits of complex I, including NDUFA4, NDUFA6, NDUFAB1, NDUFB6, NDUFB8, and NDUFB9 (representative subset shown as Fig. 6C&D). Previous work suggests that genetic ablation of PERK leads to reduction in mitochondrial supercomplex (SC) levels by blue-native PAGE^79^. To determine if there were changes in other oxidative phosphorylation complexes, we analyzed the levels of protein markers for complex II-V and found that SDHA (complex II), ATP5A (complex V), and COX11 (complex IV) levels were not altered by PERK KO (Fig. S8B & C). In conclusion, multiple subunits of complex I are PERK-dependent even in the absence of apparent ER stress.

**Figure 6:**
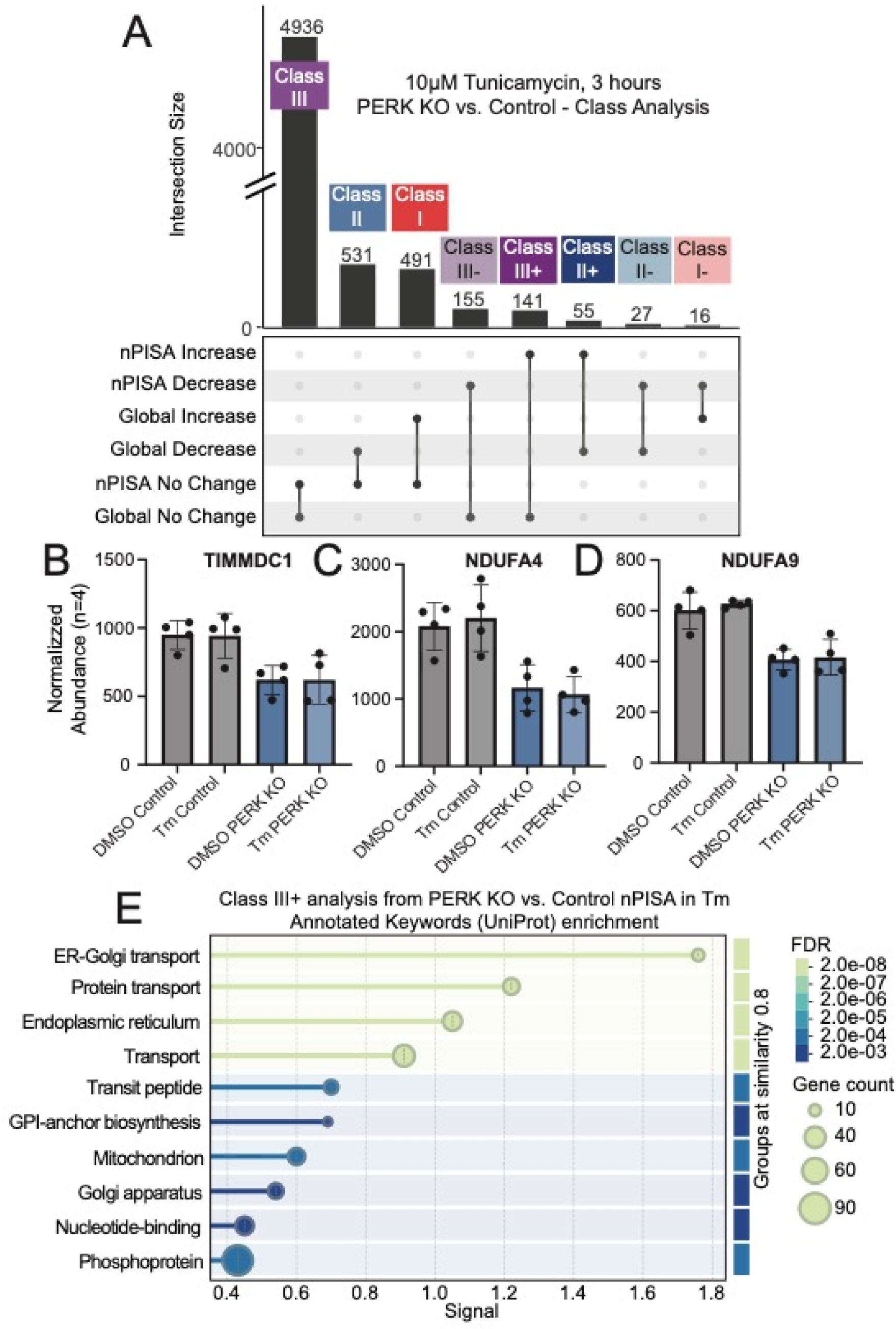
UPR signature analysis for PERK KO cells reveals proteomic changes across the endomembrane system. **A.** Upset plot of UPR Signature Class proteins in Tm treated PERK KO or WT cells. Data represent the number of intersections between Global proteome and nPISA quantitative proteomics analysis. Specific class designations are shown at the top of each bar while the graph under the bar plot shows the intersecting groups. The y-axis represents the class size with a break in the axis due to the large number of proteins in Class III. **B-D**. Global abundance in DMSO and Tunicamycin treated PERK KO or WT HEK293A cells of (B) TIMMDC1, (C) NDUFA4, (D) NDUFA9. Normalized abundance values represent the average plus or minus standard deviation (n=4). Individual biological replicate values are provided as dots. **E**. Annotated Keyword enrichment analysis from Uniprot for Class III+ proteins (n=141) defined by PERK KO vs. WT quantitative proteomics experiments from 3hr Tm treatment. Enrichment analysis was performed using String version 12.0. Keywords are grouped by similarity (0.8) with gene/protein count represented by the circle for each diagram and FDR represented by the color scale.

Multiomics analysis further identified 314 proteins that were increased in abundance in both DMSO and Tm treatment datasets. For these proteins, we applied an additional cutoff for proteins that were increasing at least 50% in protein abundance and then performed a STRING network analysis shown in Fig. S8E. KEGG analysis of these proteins found functional enrichment in p53 signaling (Fig. S8E with fold enrichment = 1.48, and FDR=2.9 e-04). P53 (TP53) was detected in our PERK KO vs. WT datasets but was not significantly changed, suggesting that the signaling changes were not necessarily due to altered TP53 levels. A key node in this network is CDKN2A which has multiple transcript isoforms which function upstream of p53^80^. SFN (14-3-3 protein sigma) and KRT17 are also both significantly upregulated with loss of PERK (Fig. S8E, Table S4). Both SFN and KRT17 have been implicated in protein synthesis regulation through Akt/mTOR signaling^81^. Several proteins involved in reduction-oxidation (REDOX) homeostasis were also identified as significantly increased proteins in PERK KO cells including HMOX1, HSP1B, NQO1, and catalase (CAT) (Fig. S8E, Table S4). These findings suggest that loss of PERK leads to protein increases in pathways related to cell growth, DNA damage, and REDOX homeostasis. Since PERK KO cells did not display any significant changes in cell growth rate (Fig. S3D), it is proposed that these protein expression changes occur to maintain cellular growth control. These findings are consistent with prior work implicating PERK in the moderation of reactive oxygen species^82^, but identify specific PERK-dependent targets for follow-up.

PERK-dependent thermal stability changes in Tm treatment were readily identified with nearly 3-times as many proteins identified as thermally altered in DMSO (compare Fig. 6A to Fig. S8A, Tables S5 and S6). In both Tm and DMSO treatment studies, the largest set of thermally altered proteins were in class III in the PERK KO vs. WT datasets. To provide biological insights into the thermal changes that were observed, we performed Uniprot keyword enrichment analysis on Class III+ proteins (Fig. 6E). PERK loss exacerbates changes in the protein thermal states specifically within the ER but also causes additional changes in mitochondrial proteins and those involved in transport between the ER and the Golgi (Fig. 6E). Of note, there was enrichment in proteins annotated as containing transit peptides, which are responsible for trafficking nuclear DNA encoded-proteins to the mitochondria^83^. Additionally, multiple proteins involved in GPI-anchor biosynthesis were thermally stabilized. Prior studies have shown that multiple GPI-anchored proteins are enriched in MAMs as are PERK and multiple other thermally stabilized proteins including Wolframin^84,85^. These results suggest that there are multiple changes in protein thermal stability mediated by PERK during 3-hour Tm treatment within and outside the endomembrane system including the ER, mitochondria-associated endoplasmic reticulum membranes, mitochondria, and Golgi. Overall, our findings show the utility of these datasets for the identification of novel PERK-dependent and -independent protein changes in the presence or absence of ER stress induction.

### nPISA is an effective system-wide tool for UPR

Our findings illustrate the utility of protein-based multiomics using nPISA and global proteomics for novel discovery. Our studies extend the traditional application of thermal shift methods such as PISA beyond the detection of small molecule or drug targets and towards analysis of system biology^2^. By its nature, the UPR would present a unique challenge to thermal shift methods because of large-scale changes in protein folding, induced protein expression, and protein degradation, along with the potential for protein aggregation events. Our studies show that nPISA effectively characterizes system-wide changes associated with drug mechanism of action even in combination with genetic perturbation in the case of PERK KO studies. Our extensive accounting of proteins targeted by ERAD following 3 hours of Tm treatment can be used as a resource for ERAD target identification and possibly substrate prioritization for degradation during the UPR.

Our protein-based multiomics analysis found that mitochondrial homeostasis is specifically altered in the absence of PERK and is reflected in changes in thermal stability and protein abundance. The role of PERK in mitochondrial regulation continues to emerge with strong evidence suggesting that MAM dynamics play a significant role in mitochondrial health. Our studies revealed changes specifically in complex I subunit abundance which was accompanied by multiple changes in multiple REDOX related factors. Additionally, our thermal stability studies identified enrichment in not only proteins that are resident to the ER, but also in proteins containing transit peptides suggesting that PERK could influence targeting of nuclear encoded mitochondrial factors in a manner that alters their biophysical state. From our studies, we cannot rule out that transit peptide containing-proteins are altered because of the altered redox dynamics associated with PERK loss rather than by direct PERK regulation.

Finally, we provide an extensive data-driven multiomics class analysis that identifies novel protein class responses to Tm-induced UPR regardless of functional annotation status. This approach represents a novel path to protein functional annotation based on multiple protein features which can then be compared to existing functional categories as a reference. Thermal shift analyses such as nPISA have great potential for functional annotation because many different changes can alter protein thermal stability by nPISA including changes in protein-protein or other protein interactions, protein post-translational modification, protein sequence variant presence, and protein folding status. Protein-based multiomics with post-translational modification (PTM) mapping, such as phosphoproteomics, has been extensively used for system characterization but suffers from a wide-spread lack of annotation for functional modification sites. As a consequence, protein thermal stability analyses such as nPISA will likely outperform PTM studies, as shown here, since many PTMs do not alter protein function. The complexity of UPR signaling, characterized by diverse protein modifications, dynamic folding states, and the essential role of ERAD in apoptosis prevention, makes the UPR a compelling model to showcase nPISA as a foundational approach for functional proteomics

## Supplemental Tables

**Table S1**. **Global and nPISA analysis for parental HEK293A cells treated with DMSO or Tunicamycin for 3 hours.** Table contains global abundance and normalized PISA values for the DMSO vs. Tunicamycin datasets (n=4 per treatment) including “Abundance Ratio”, “Abundance Ratio p-value”, “Abundance Ratio adjusted p-value”, Protein Uniprot Accession numbers, Protein Descriptions, Number of Peptide Spectrum Matches (PSMs), and Number of Unique Peptides. Global Data and nPISA Data are presented in separate columns designated at the top of the sheet in red font.

**Table S2**. **Global and nPISA analysis for clonal PERK KO and WT cells treated with DMSO or Tunicamycin for 3 hours.** All quantitative proteomics comparisons for the comparison of Tunicamycin and DMSO treatment in the genotypes as stated above. Specific tabs include tables for nPISA calculations for Tm treatment in PERK KO vs. WT cells, nPISA calculations for DMSO treatment in PERK KO vs. WT cells, global abundance analysis for Tm treatment in PERK KO vs. WT cells, global abundance analysis for PERK KO in either Tm or DMSO treatment, global abundance analysis for DMSO treatment in PERK KO vs. WT cells, global abundance analysis for WT cells in either Tm or DMSO treatment. Columns for each table include Uniprot Accession number, Protein descriptions, Average Ratios between the treatments designated in each tab, p-values for those comparisons, log2 calculations for the abundance values, and –log10 (p-value) calculations for each comparison.

**Table S3**. **Protein multiomics class analysis of HEK293A cells with Tm vs. DMSO treatment**. Uniprot accession numbers for each protein associated with a given protein-based multiomics analysis class as described in Figure 5.

**Table S4**. **Significantly altered protein change intersection between clonal PERK KO and WT cells with DMSO or Tm treatment in global proteomics.** Table contains global abundance values for the DMSO vs. Tunicamycin datasets (n=4 per treatment) in PERK KO vs. WT cells for all changes with p-values >0.05 in both treatments defining PERK regulated proteins that are not ER stress dependent. The table includes column headers “Abundance Ratio” for each comparison, “Abundance Ratio p-value” for each comparison, “Abundance Ratio adjusted p-value” for each comparison, Protein Uniprot Accession numbers, Protein Descriptions, “Coverage”, Number of Peptide Spectrum Matches (PSMs), and Number of Unique Peptides.

**Table S5**. **Protein multiomics class analysis of clonal PERK KO and WT cells with Tm treatment.** Table includes all proteins associated with a given protein-based multiomics analysis class for comparison of PERK KO vs. WT cell global and PISA analysis in Tunicamycin treatment conditions. Classes are designated in tabs with “+” is represented by “P” and “-“ is represented by “M”. This includes the Uniprot Accessions, Protein Descriptions, FC_global_Tm (FC=fold change), pvalue_global_Tm, FC_nPISA_Tm, and pvalue_nPISA_Tm.

**Table S6. Protein multiomics class analysis of clonal PERK KO and WT cells with DMSO treatment.** Table includes all proteins associated with a given protein-based multiomics analysis class for comparison of PERK KO vs. WT cell global and PISA analysis in DMSO treatment conditions. Classes are designated in tabs with “+” is represented by “P” and “-“ is represented by “M”. This includes the Uniprot Accessions, Protein Descriptions, FC_global_Tm (FC=fold change), pvalue_global_Tm, FC_nPISA_Tm, and pvalue_nPISA_Tm.

## Supplemental Figures

**Figure S1.**
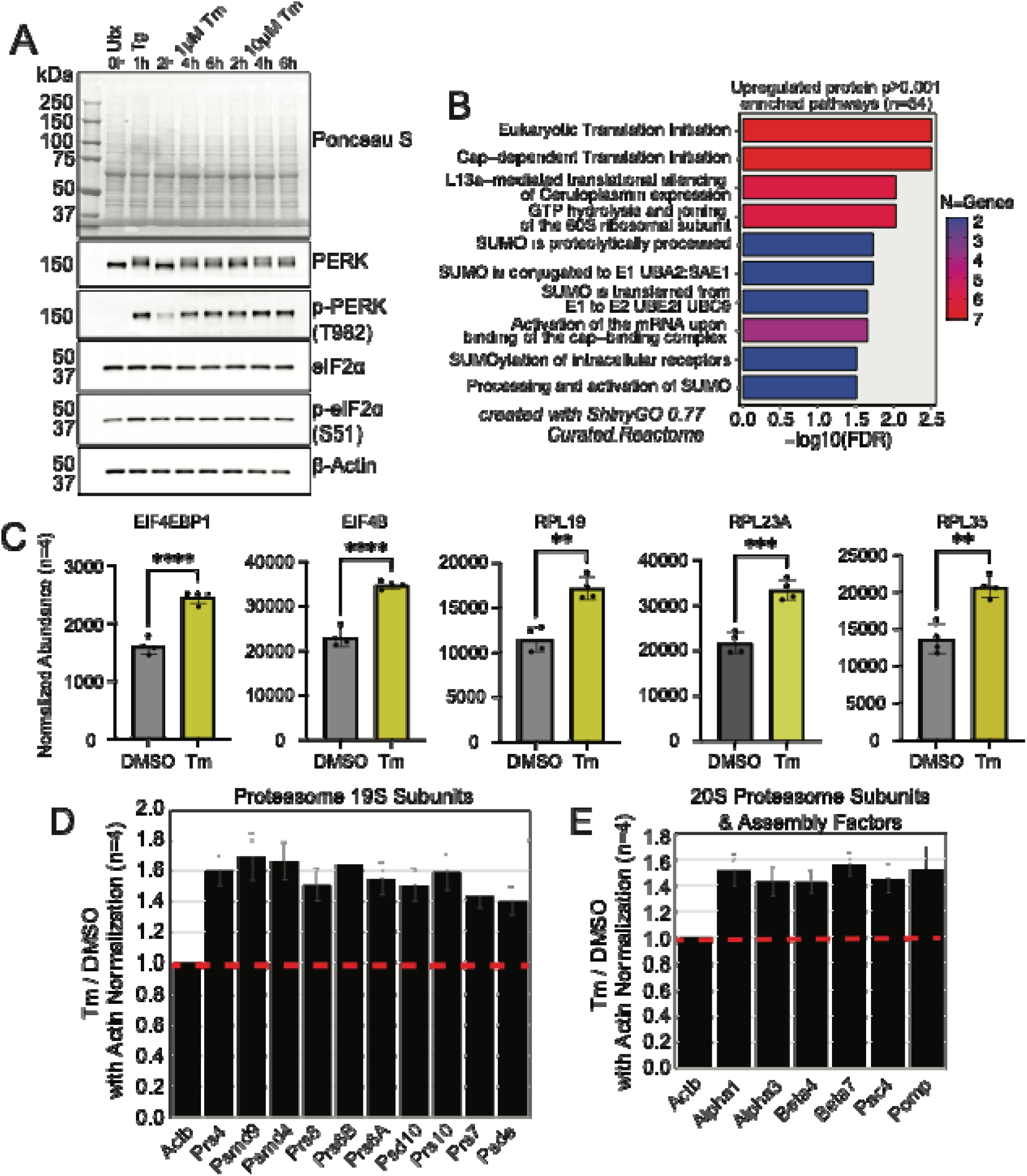
Global proteome analysis identifies upregulation of proteins involved in translation and protein degradation. **A**. Comparison of 1 μM thapsigargin or 10 μM tunicamycin treatment of HEK293A measured by immunoblot of UPR proteins. Time course experiment from 1-6 hours. Ponceau S analysis along with immunoblot for UPR markers include PERK kinase, phospho-PERK (T982, p-PERK) eIF2α, phospho-eIF2α (S51, p-eIF2α), and β-Actin as a loading control. **B**. ShinyGO enrichment analysis of upregulated proteins with an adjusted p-value ≤ 0.05 (n=54) using Reactome. Data are represented as a bar graph of the -log10 FDR values for the enriched pathways. The enriched pathways are listed to the left and a color scale representing the number of genes/proteins in each category is shown to the right. **C, D**. Quantitative proteomics data showing increased abundance of multiple subunits of the 19S proteasome (C) or 20S proteasome (D) scaled to the normalized abundance values to β-Actin (n=4). The red line indicates no change in abundance between DMSO and Tm samples.

**Figure S2:**
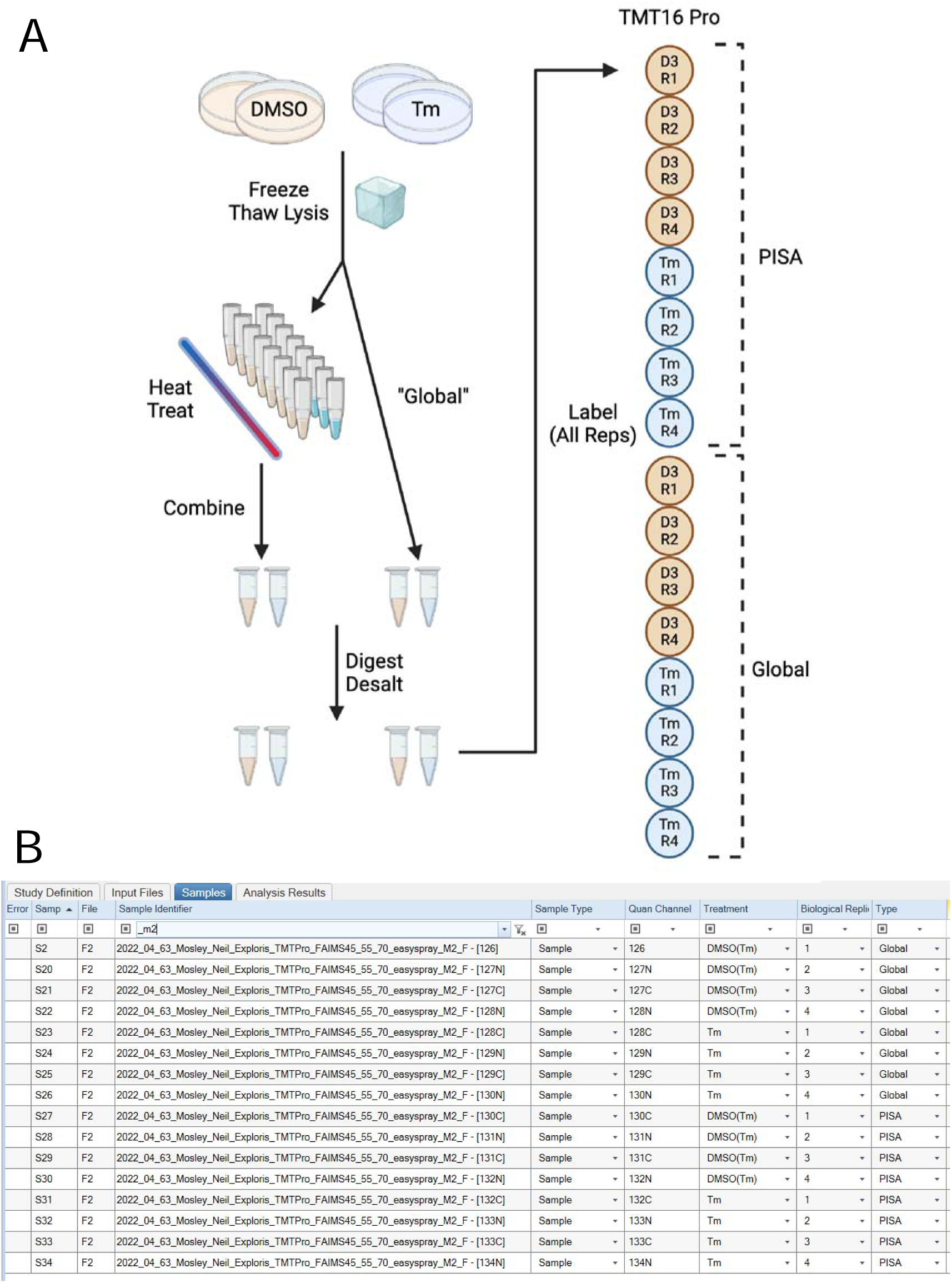
Multiplex design for global proteomics and PISA experiments comparing Tm treatment versus DMSO. **A**. Schematic representation of the PISA (see heat treatment) and global proteomics experiments. Individual biological replicates were prepared and labeled with TMT16Pro **B**. Details of Proteome Discoverer 2.5 Samples that illustrate the TMT channel used for each experiment and biological replicates for this study.

**Figure S3.**
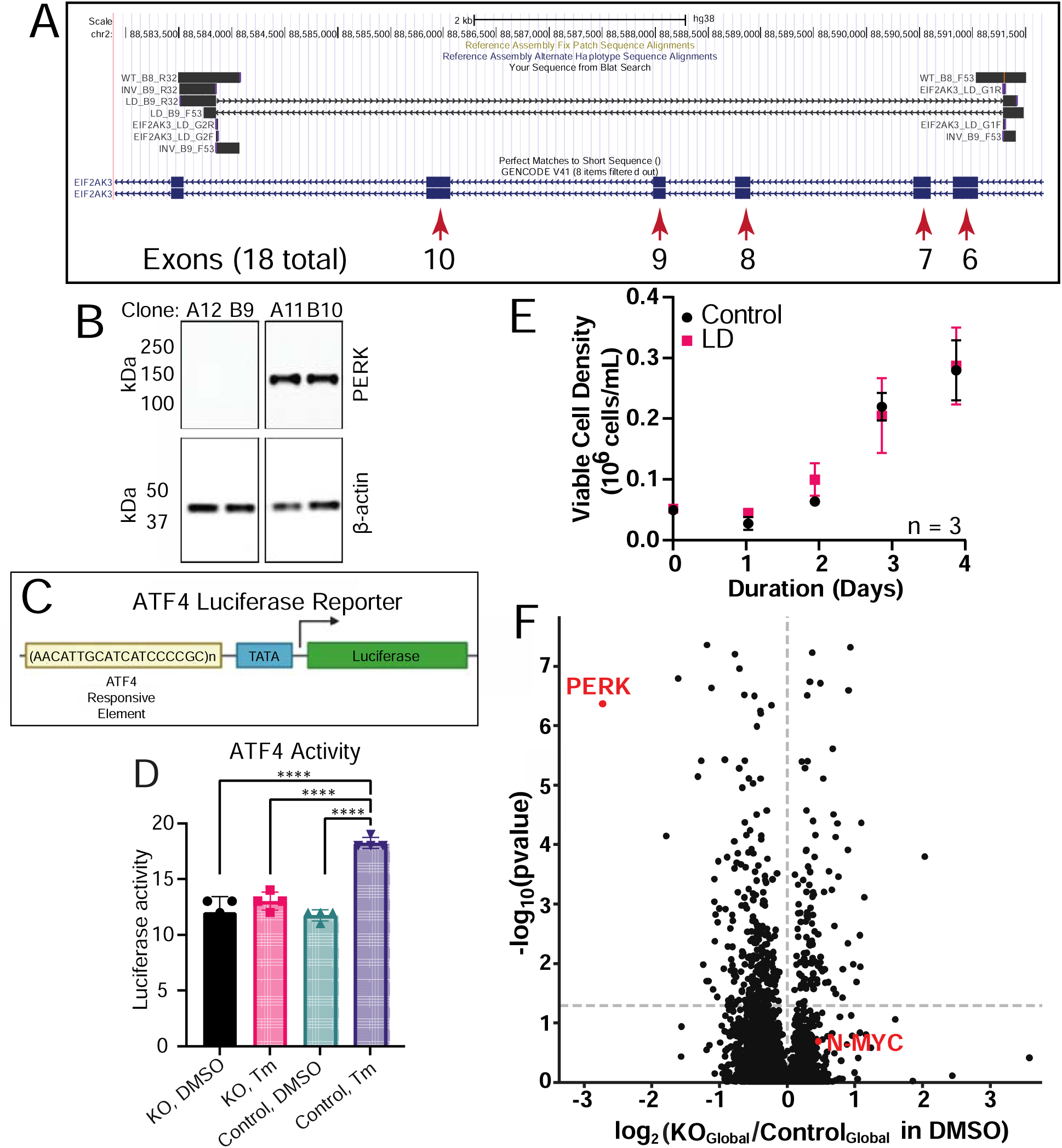
Characterization of clonal PERK KO HEK293A cell lines. **A**. Alignment of EIF2AK3 locus with regions amplified in clonal cell lines by R32 and F53 primers. Confirmation of PERK KO shown for clone B9 used in the proteomics studies. View is zoomed into exons affected by CRISPR-Cas9 deletion. **B**. Immunoblot analyses of PERK KO (clones A12 and B9 shown) and WT (clones A11 and B10 shown) single clonal cell lines. Lysates were prepared and PERK and β-actin were measured using specific antibodies by immunoblot analyses. **C**. Luciferase reporter assay for measuring ATF4 transcriptional activity in HEK293A cell lines. **D**. Measurement of ATF4-transcriptional activity was measured using a stably expressed luciferase reporter in WT HEK293A cells and PERK KO following either DMSO or Tm treatment. Results are shown in a bar graph +/- standard deviation, with each replicate values shown as a dot (n=4). **E.** Growth analyses of WT versus PERK KO clonal cell lines. Points shown are average cell density values +/- standard deviation (n=3) over a 4-day time course. **F**. Volcano plot showing changes in global protein abundance as measured in WT and PERK KO cells treated with DMSO (n=4). The y-axis shows the -log10p-value for each protein measurement (represented by a dot). Red dots are shown for quantitation of PERK kinase and the N-Myc transcription factor. The dashed line on the figure represents a p-value ≤ 0.05.

**Figure S4:**
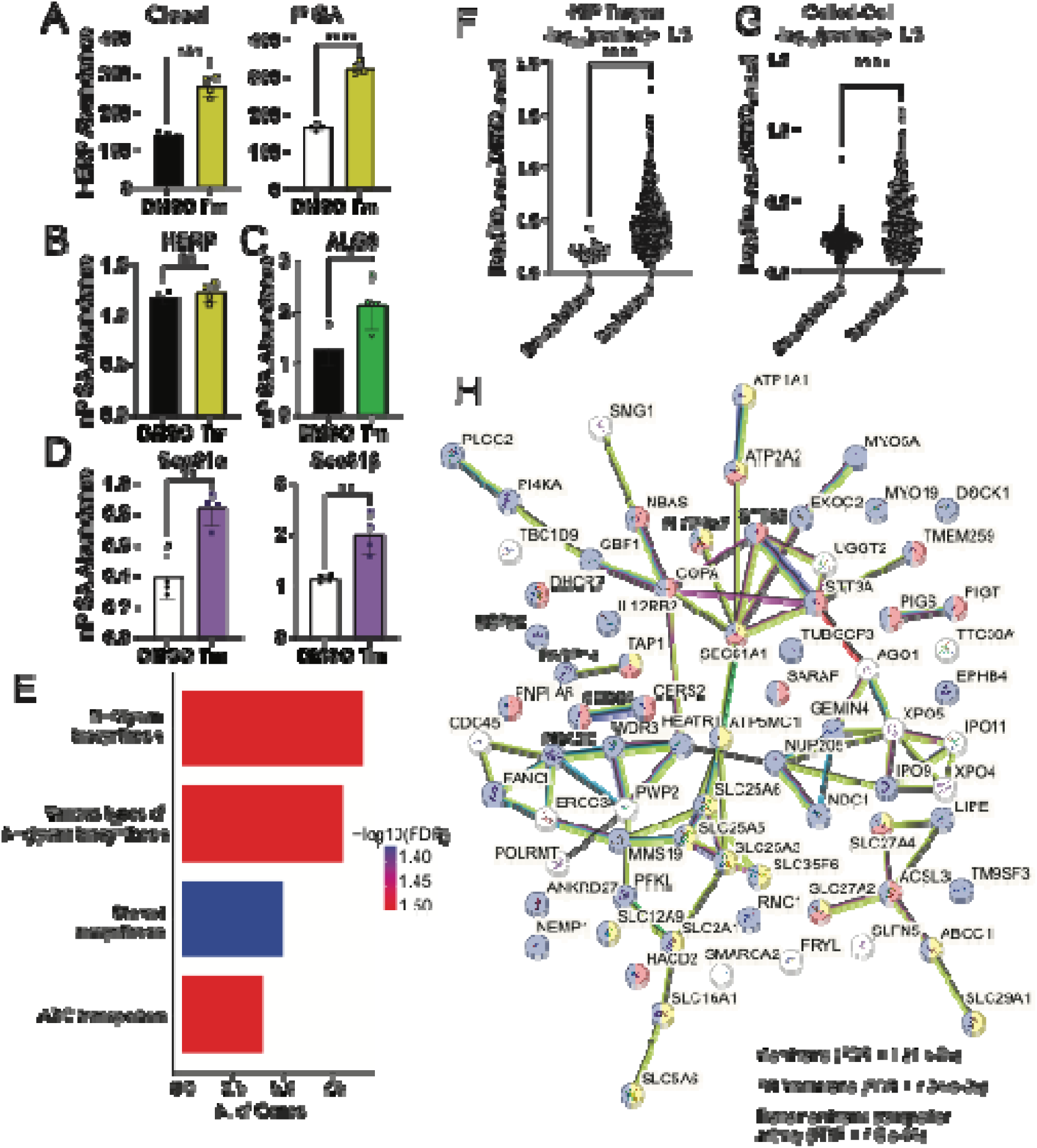
nPISA reveals thermal stability changes in ER-associated and heat shock pathways. **A**. Global and PISA data for the UPR-induced protein HERP show increases related to gene expression induction. Abundance values from the different quantitative experiments as indicated at the top of each graph in either DMSO or Tm treatment conditions as indicated. Data are shown as averages plus or minus standard deviation with dots for each replicate value shown (n=4). **B**. Normalization of PISA data by global protein abundance (i.e. the input value) corrects false identification of thermal stability changes for differentially abundant proteins. Data are shown as in A. **C**. ALG9 nPISA abundance levels shown as in A. **D**. Sec61α and β nPISA abundance levels shown as in A. **E**. ShinyGO analysis of Reactome pathway enrichment for proteins that were significantly thermally altered according to nPISA (n=4) with an adjusted p-value ≤ 0.05. Bars show the total number of genes/proteins within each enrichment term with color coding based on the -log10 of the FDR. Protein background used for this enrichment analysis was an input of all proteins quantified by LC-MS/MS for the Tm vs. DMSO experiments. **F&G**. Absolute value plot showing the data distribution for proteins that (F) were identified as HSP targets or (G) are annotated as Coiled-Coil domain containing that were significantly destabilized or stabilized by Tm treatment according to nPISA measurement. **H**. STRING (version 12) network analysis of the proteins shown within the red box in Figure 3F. Nodes are color coded as indicated in the key below the network, with clear nodes indicating those that are not annotated within the designated functional categories.

**Figure S5.**
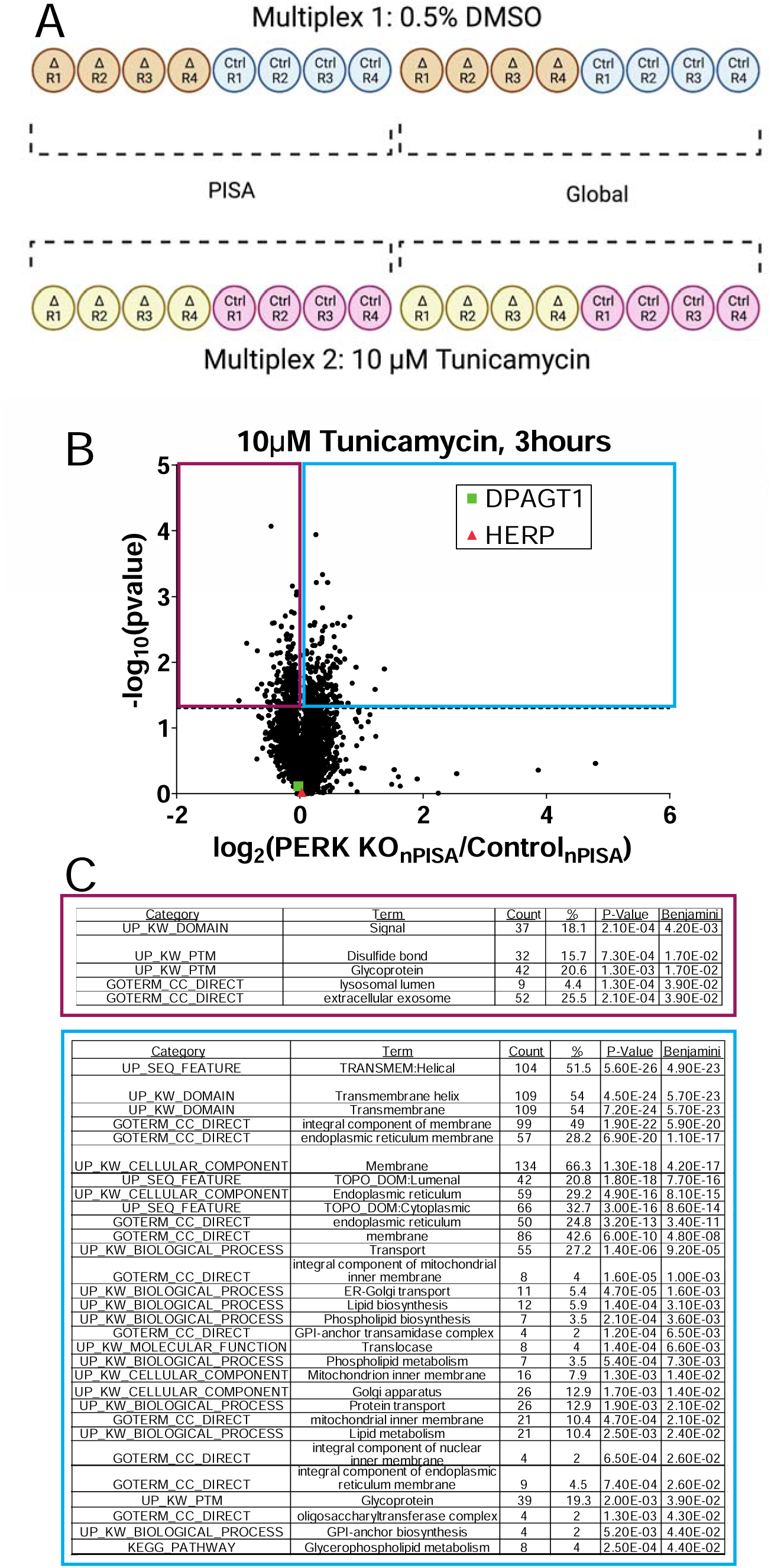
nPISA Analysis of PERK KO and WT cell lines reveals PERK-dependent protein stability changes. **A.** TMT multiplex design for PERK KO vs. WT HEK293A cells. An additional multiplex was performed for the global proteomics samples for both sets. Data from these studies is provided as Table S2. **B.** Volcano plot of nPISA data from PERK KO and WT cells treated with Tunicamycin with DPAGT1 (target of Tm) and HERP (ERAD protein) highlighted. Dots represent the average log2 nPISA ratio between PERK KO and WT cells. The purple box designates proteins that are significantly destabilized in the PERK KO with a p-value ≤ 0.05 whereas the blue box designates proteins that are significantly stabilized in the PERK KO using the same statistical cutoff. **C**. Database for Annotation, Visualization, and Integrated Discovery (DAVID) analysis of proteins in the destabilized and stabilized groups as described in B. The protein background for these studies was set as all proteins that were quantified in the Tunicamycin multiplex shown in A. Statistically enriched functional groups are shown that had a Benjamini correct p-value cutoff ≤ 0.05. Thermally stabilized proteins are shown in purple and destabilized in blue.

**Figure S6:**
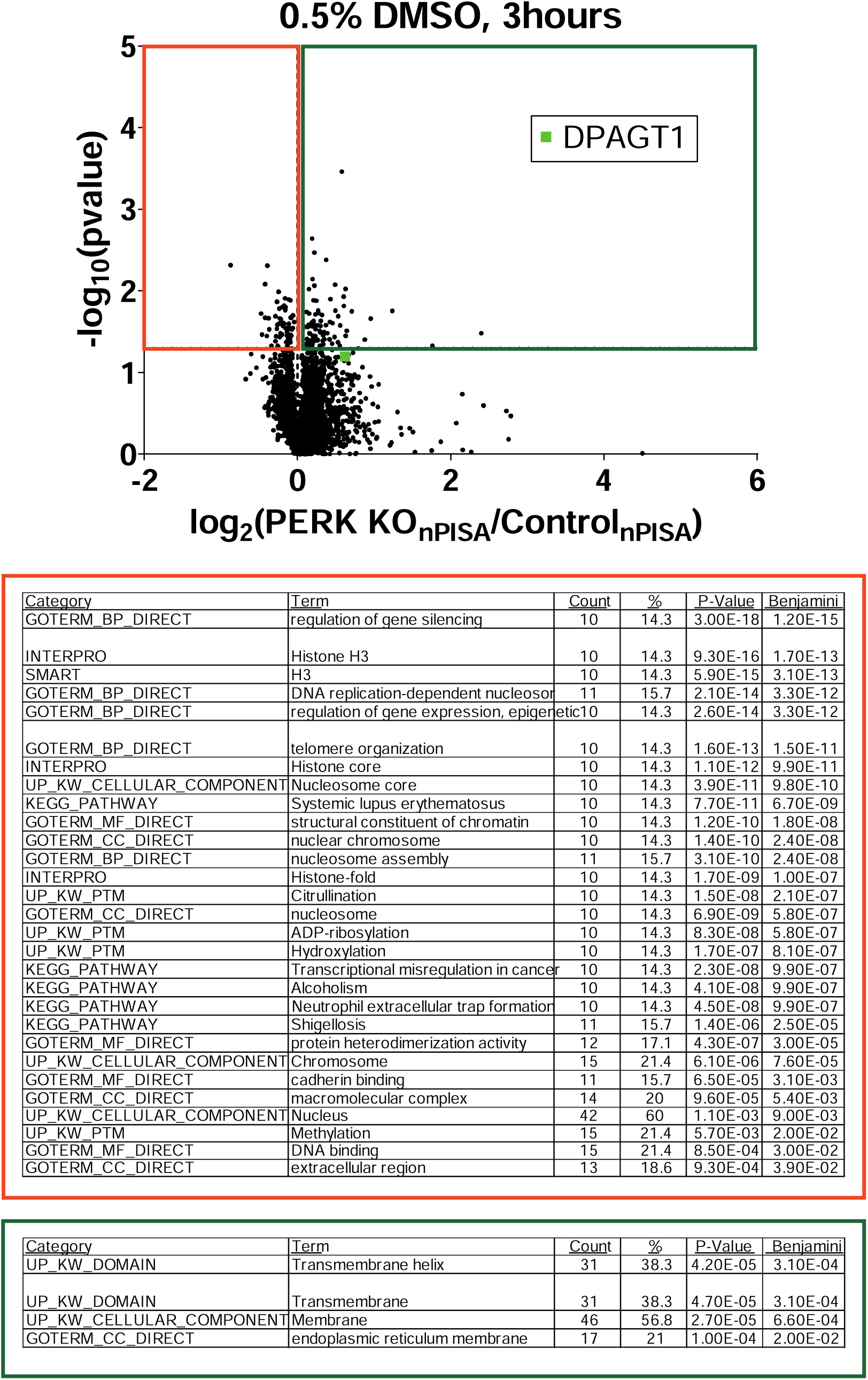
nPISA and functional enrichment analysis of PERK KO and WT Cell Lines with DMSO treatment. **A**. Volcano plot of nPISA data from WT and PERK KO cells treated with DMSO for 3hrs. Dots represent the average log2 nPISA ratio WT/PERK KO cells. The red box designates proteins that are significantly destabilized in the PERK KO with a p-value ≤ 0.05 whereas the green box designates those that are significantly stabilized in the PERK KO using the same statistical cutoff. DPAGT1 (green) is a known target of Tm, along wth HERP1 (red). A lower number of statistically significant nPISA changes was observed with DMSO treatment relative to Tm (Fig. S5). **B**. DAVID analysis of proteins in the destabilized and stabilized groups as described in A. The protein background for these studies was set as all proteins that were quantified in the multiplex shown in A. Statistically enriched functional groups are shown that had a Benjamini correct p-value ≤ 0.05. The categories in the red box represent thermally destabilized protein enrichments whereas the green box represents thermally stabilized protein enrichment groups.

**Figure S7:**
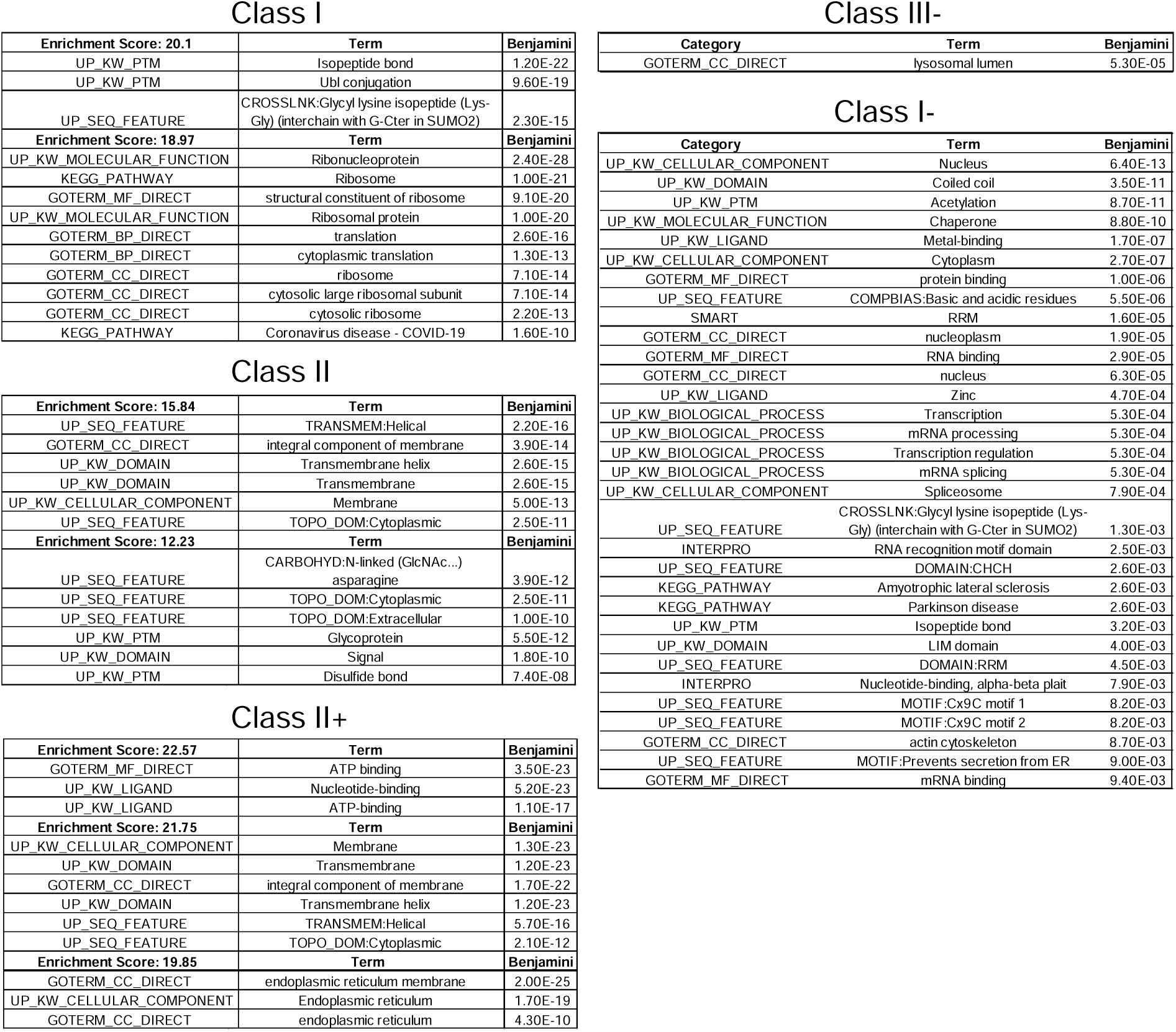
Summary of DAVID bioinformatic analyses for MultiOMICS interaction classes. Classes are described in Figure 6 from the Tm vs. DMSO treatment studies of parental HEK293A PISA experiments. Note that not all classes resulted in significant pathway enrichments and in those cases no data is shown. Background data used for the enrichment studies was customized using only proteins that were quantified in the Tm vs. DMSO treatment studies (Table S1).

**Figure S8:**
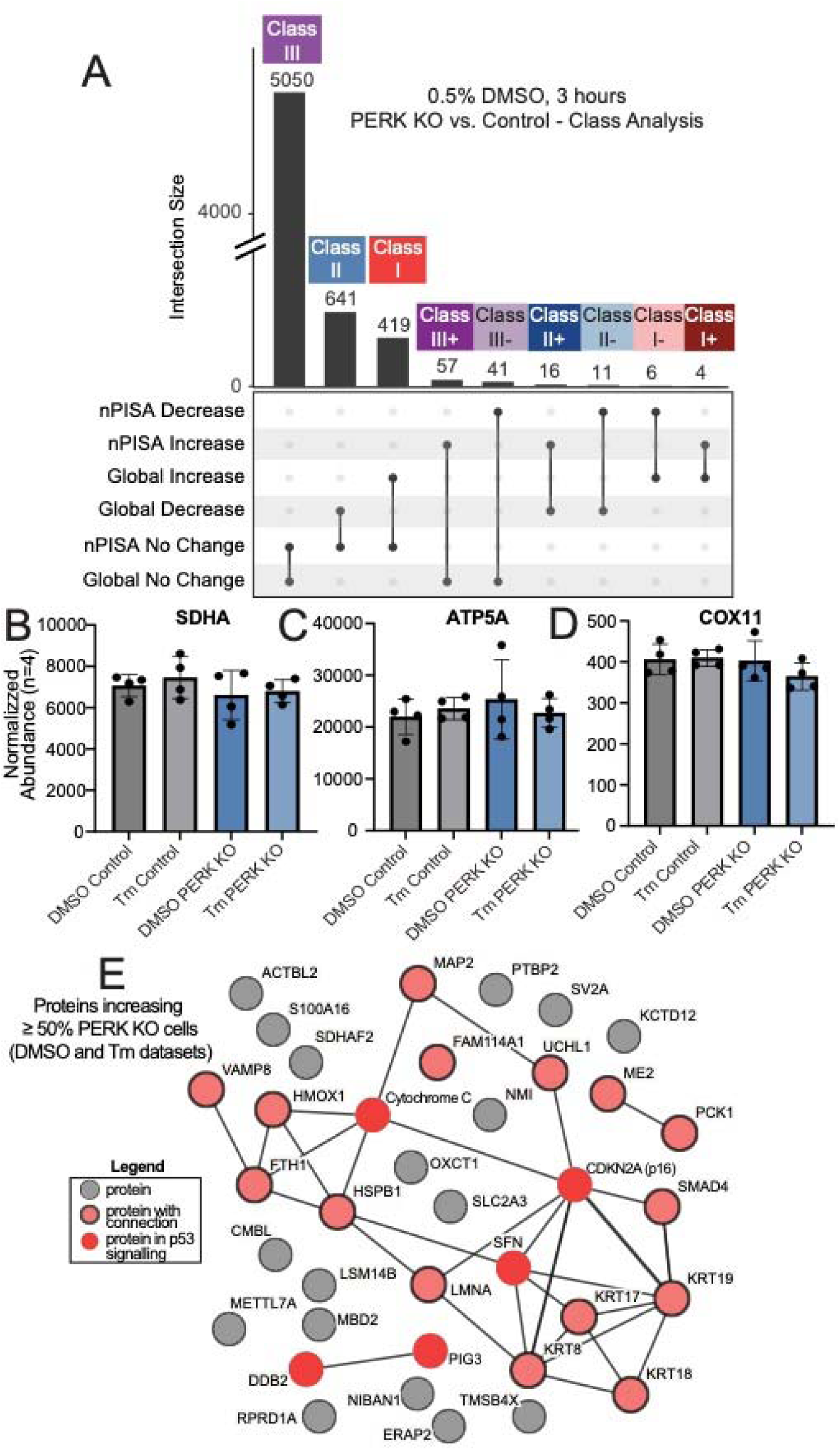
Class analysis of WT versus PERK KO cells using multiOMICS. **A.** Upset plot illustrating overlap and designation of UPR Signature Class proteins in DMSO Global proteome and nPISA quantitative proteomics analysis. Each bar represents the total number of proteins in each class. For this analysis, proteins that were not identified in both the global and nPISA studies are not included. Specific class designations are shown at the top of each bar while the graph under the bar plot shows the intersecting groups for reference. The y-axis represents the class size with a break in the axis due to the large number of proteins in Class III. **B-D**. Global abundance in DMSO and Tm treated PERK KO or WT HEK293A cells of mitochondrial complex subunits (B) SDHA, (C) ATP5A, and (D) COX11. Normalized abundance values represent the average plus or minus standard deviation with individual biological replicate values provided as dots (n=4). Protein abundance levels were determined to not be statistically different by t-test (n=4). **E**. STRING version 12 network analysis of proteins significantly increasing at least 50% in global abundance in both the Tm and DMSO datasets for PERK KO vs. WT suggesting strong expression dependent on PERK. A legend describing the color coding for each node (circles) is shown to the left.

## Acknowledgements

A portion of the funding for this project was provided by the T32DK064466 Diabetes and Obesity Research Training Program at IUSM, DeVault Fellowship (N.M.), the Pediatric and Adult Translational Cancer Drug Discovery and Development Training Program T32CA272370 (A.R.) and F30AG079580 from NIA (H.R.S.W). Additional support was provided by the Showalter Research Trust (A.M.) and by the Indiana University Melvin and Bren Simon Cancer Center (P30CA082709). Partial support for this project was provided by NIH (to RCW: GM136331; to ALM: NS121550 and CA082709) and by the Indiana Clinical and Translational Sciences Institute, Award Number UL1TR002529. The content is solely the responsibility of the authors and does not necessarily represent the official views of funders.

## Author Contributions

Experiments were designed by N.A.M., A.L.M., and R.C.W, Experiments were executed by N.A.M., A.L.M, W.R.S.K and E.D. Cell line development was managed by S.P. and N.A.M. Data analysis was performed by N.A.M., A.L.M, H.R.S.W., A.M.R., and S.A.P.J. Manuscript was written by N.A.M., A.L.M, S.A.P.J., and R.C.W.. Experiments were supervised by A.L.M. and R.C.W.

## Declaration of Interests

R.C.W. is a scientific advisor to HiberCell. The other authors declare no competing interests.

## Experimental Procedures

### Cell Culture and ER stress

The study used HEK293A cells (Invitrogen, Carlsbad, CA) transduced with a lentivirus encoding an ATF4 transcription reporter (P(AAREx6)-Luc), which consists of six copies of C/EBP-ATF4 consensus binding sequence upstream of a minimal promoter and the firefly luciferase coding sequence ^28^. Cells were cultured Corning Dulbecco’s Modified Eagle Medium (DMEM, 10-013CV) supplemented with 10% fetal bovine serum (FBS) at 37°C with 5% CO_2_ and humidification. Cultures were passaged every 3-4 days to maintain viability and were also periodically checked for absence of mycoplasma contamination. Treatment experiments consisted of removing growth medium and replacing with fresh media supplemented with tunicamycin (Calbiochem 654380), or DMSO. After culturing for 3hrs, the media was aspirated from the plates and cells were washed with 1X PBS prior to either lysis or removal from the plates.

### SDS-PAGE and Immunoblotting

Lysates were prepared for immunoblot experiments by removing cells from the plates with a rubber scraper. Proteins were separate by SDS-PAGE using 4% stacking and 7.5% resolving gels and transferred to filters that were incubated with antibodies specific to SERCA2A (Abcam ab2861, lot GR3275540-3), PERK (Cell Signaling C33E10, Lot 11), p-PERK (p-PERK-T982, Custom rabbit monoclonal rIgG; LLY-71, clone 1-6, Lot #BE02734-048), Beta-Actin (Sigma A5411-2ML, lot 0000120485), ATF4 (In-house antibody), eIF2α (Cell Signaling 5324S, Lot 7), p-eIF2α (abcam ab32157, lot GR3254220-12), Ire1 (Invitrogen MA5-14991, Lot XH3669534), BiP (HSPA5/GRP78) (Invitrogen MA5-27686, Lot XH3667743), XBP1-s (Cell Signaling E9V3E, 40435S, Lot 2). SERCA2A antibody was validated using lentivirus-based overexpression of SERCA2A. The ATF4 antibody was validated based on results from ATF4 reporter cell line. The PERK antibody was validated based on results from PERK KO cell line. The GRP78 and Ire1 antibodies both have Invitrogen “Advanced Verification” status indicating that they have been tested using knockdown or cell treatment. HRP was quantitated using Biorad Clarity Western ECL Substrate and blots were imaged using a Biorad Chemidoc. Quantitation was done using the Biorad ImageLab software. Beta-Actin was used as a loading control and figures show a representative set of Beta-Actin blots since samples were run on multiple gels.

### Proteome Integral Solubility Alteration (PISA) Experiments

Following treatment, cells were rinsed with cold PBS and harvested by scraping and centrifugation at 300 x g for 5 minutes. Cells were washed again with cold PBS, collected by centrifugation at 300 x g for 5 minutes, and flash frozen at -80 °C. Prior to execution of the TPP workflow, cell pellets were removed from the freezer and were then re-suspended in lysis buffer solution (40 mM HEPES [pH 7.5], 200 mM NaCl, 5 mM Beta glycerophosphate, 0.1 mM sodium orthovanadate, 2 mM TCEP, 0.4% NP40, and 1X Roche EDTA free mini complete protease inhibitor). Cells were snap frozen in liquid nitrogen and thawed in ice water. Lysed cells were clarified by centrifugation at 20,000 rcf for 30 minutes at 4°C. Protein concentrations were measured in supernatants by a Lowry based method and adjusted to a common concentration within each biological replicate (ca 3 mg/mL) using the lysis buffer solution. A total of 50-100ug of protein was reserved from paired samples for global proteomics studies. Aliquots of the adjusted supernatants (50 uL) for PISA studies were transferred to PCR tubes after which the samples were heated and cooled using a gradient procedure in a thermocycler. The heat treatment consisted of 2 minutes at 25°C, 3 minutes at temperature per gradient, 2 minutes at 25°C, followed by 4°C. Gradient temperatures consisted of 39.9, 43.5, 45.8, 48.4, 51.1, 53.8, 56.2, and 60.1°C. Heat treated samples were clarified by centrifugation at 20,000 rcf for 20-30 minutes to pellet precipitated protein after which the supernatant was decanted and frozen.

### Sample Preparation for LC-MS/MS and Mass Spectrometry

Supernatant samples (post heat treat and global) were first precipitated in 20% Trichloroacetic acid and washed with 100% acetone. Dried pellets were resuspended in 8M Urea in a solution of 100 mM Tris (pH 8.5). Samples were reduced with TCEP and alkylated with chloroacetamide. Reduced and alkylated samples were digested with Trypsin/LysC followed by quenching with formic acid as previously described^16^. Quenched samples were desalted with Waters C18 columns and then isobarically labeled with Thermo Scientific TMTPro labels. Raw MS files and supplemental MS data can be found for the PISA and PERK KO PISA experiments are available upon reasonable request.

The global proteomics and PISA were analyzed on an Exploris 480 mass spectrometer (Thermo Scientific) coupled to an EASY-nLC HPLC system (Thermo Scientific). The peptides were eluted using a mobile phase (MP) gradient for 180 minutes. FAIMS source with a compensation voltage of -45, -55, -70V. During peptide elution, the mass spectrometer method was operated in positive ion mode for 180 mins, programmed to select the most intense ions from the full MS scan using a top speed method. Exploris MS1 parameters include: Microscans 1; MS1 Resolution 60k; normalized automatic gain control (AGC) target 300%; and Scan range 375 to 1500 m/z. Exploris data dependent MS/MS parameters include: Microscans 1; Resolution 45k; AGC target 200%; Isolation window 0.7 m/z; Fixed first mass 100 m/z; and HCD normalized collision energy 32.0. The data were recorded using Thermo Xcalibur software (Thermo Fisher).

The samples from the global and PISA experiments were also analyzed on an Orbitrap Eclipse mass spectrometer (Thermo Scientific) coupled to an EASY-nLC HPLC system (Thermo Scientific). The peptides were eluted using a mobile phase (MP) gradient for 180 minutes. FAIMS source with a compensation voltage of -45, -55, -65V. During peptide elution, the mass spectrometer method was operated in positive ion mode for 180 mins, programmed to select the most intense ions from the full MS scan using a top speed method. Exploris MS1 parameters include: Microscans 1; MS1 Resolution 120k; automatic gain control (AGC) target 4E5; and Scan range 400 to 1600 m/z. Exploris data dependent MS/MS parameters include: Microscans 1; Resolution 50k; AGC target 1E5; Isolation window 0.7 m/z; Fixed first mass 100 m/z; and HCD normalized collision energy 34.0. The data were recorded using Thermo Xcalibur software (Thermo Fisher).

### Data Analyses

Global abundance values from mass spectrometry studies were based on reporter ion abundance and were averaged across all biological replicates for the described studies (n=4). For global abundance analysis, p-values and adjusted p-values were calculated using Proteome Discoverer using quantitative values from unique and razor peptides. PISA experiments were analyzed by first combining protein abundance across instrument technical replicates. The averaged abundances were then normalized for each biological replicate by dividing the PISA abundance for each condition (i.e. drug, cell line) by its corresponding global abundance. These normalized values (nPISA) for each biological replicate and each treatment were then used to conduct a two-tailed, two-sample equal variance t-test along with corresponding -log_10_(t-test) to calculate the statistical significance of each shift (p-value) with post-hoc adjustment of the p-values for nPISA performed using Bonferroni Correction. The nPISA values for each biological replicate were also averaged for each condition (i.e. PERK KO) after which the ratio of treatment to WT (i.e. nPISA_KO_/nPISA_WT_) was determined and used for fold change calculation (i.e. log2(nPISA_KO_/nPISA_WT_)) and significance calculations performed as described above. Multiomics analysis was performed using intersection analysis using R and included use of UpSetR ^86^, Clustvis ^87^, and ggplot2. Data visualization was also performed using GraphPad Prism.

